# Energy landscape analysis elucidates the multistability of ecological communities across environmental gradients

**DOI:** 10.1101/709956

**Authors:** Kenta Suzuki, Shinji Nakaoka, Shinji Fukuda, Hiroshi Masuya

**Affiliations:** Integrated Bioresource Information Division, BioResource Research Center, RIKEN, 3-1-1 Koyadai, Tsukuba, Ibaraki 305-0074, Japan; Laboratory of Mathematical Biology, Faculty of Advanced Life Science, Hokkaido University, Kita-10 Nishi-8, Kita-ku, Sapporo, Hokkaido 060-0819, Japan; PRESTO, Japan Science and Technology Agency, 4-1-8 Honcho, Kawaguchi, Saitama 332-0012, Japan; Institute for Advanced Biosciences, Keio University, 246-2 Mizukami, Kakuganji, Tsuruoka, Yamagata 997-0052, Japan; Intestinal Microbiota Project, Kanagawa Institute of Industrial Science and Technology, 3-25-13 Tonomachi, Kawasaki-ku, Kawasaki 210-0821, Kanagawa, Japan; Transborder Medical Research Center, University of Tsukuba, 1-1-1 Tennodai, Tsukuba, Ibaraki 305-8575, Japan

**Keywords:** community assembly, compositional stability, historical contingency, multistability, alternative stable states, regime shift, presence/absence data, Markov network, pairwise maximum entropy model, principle of maximum entropy, stability landscape, energy landscape

## Abstract

Compositional multistability is widely observed in multispecies ecological communities. Since differences in community composition often lead to differences in community function, understanding compositional multistability is essential to comprehend the role of biodiversity in maintaining ecosystems. In community assembly studies, it has long been recognized that the order and timing of species migration and extinction influence structure and function of communities. The study of multistability in ecology has focused on the change in dynamical stability across environmental gradients, and was developed mainly for low-dimensional systems. As a result, methodologies for studying the compositional stability of empirical multispecies communities are not well developed. Here, we show that models previously used in ecology can be analyzed from a new perspective - the energy landscape - to unveil compositional stability in observational data. To show that our method can be applicable to real-world ecological communities, we simulated assembly dynamics driven by population level processes, and show that results were mostly robust to different simulation assumptions. Our method reliably captured the change in the overall compositional stability of multispecies communities over environmental change, and indicated a small fraction of community compositions that may be channels for transitions between stable states. When applied to murine gut microbiota, our method showed the presence of two alternative states whose relationship changes with age, and suggested mechanisms by which aging affects the compositional stability of the murine gut microbiota. Our method provides a practical tool to study the compositional stability of communities in a changing world, and will facilitate empirical studies that integrate the concept of multistability from different fields.

## Introduction

The order and timing of species migration and extinction during community assembly influence the structure (Drake 1991, Fukami and Morin 2003, Kadowaki et al. 2012) and function (Fukami et al. 2010, Jiang et al. 2011) of communities, resulting in multistability (also known as alternative stable states) of different community compositions (Fukami 2010, Fukami 2015). Compositional dynamics play a prominent role in real world ecosystem organization, and understanding community assembly in terms of the management of ecological systems has direct relevance to conservation biology, agriculture, and medicine (Fukami et al. 2015). There is a need for development of a methodology that accounts for compositional stability in multispecies communities to facilitate predicting, preventing and controlling large-scale shifts in community compositions.

In the field of microbial ecology, the recent development of next-generation sequencing technology has made it possible to comprehensively study community structures (Caporaso et al. 2010, Ding and Schloss 2014, Thompson et al. 2017, Bolyen et al. 2019). This development led to recognition of the association between composition and function of microbial communities that is essential for the maintenance of host organisms or physical environment (Costello et al. 2012, Widder et al. 2016, Sommar et al. 2017).

In the animal intestine there is a phenomenon called dysbiosis in which function is severely impaired due to infection or other causes (Carding et al. 2015). One example would be *C. difficile* infection (CDI) (Kelly and LaMont 2008, Britton and Young 2014) that can occur when the gut microbiota has been disrupted from its normal balance, e.g., by antibiotics. The normal microbiota is resistant to *C. difficile* colonization, whereas it is significantly altered when *C. difficile* successfully colonizes the intestine. There would be at least two community compositions (with or without *C. difficile*) that are stable at the same environmental conditions. In this case, transplantation of the normal microbiota can effectively restore the infected microbiota back to normal (Bakken et al. 2011). However, simply implanting a desirable microbiota is not a universal method for its establishment (Rilling et al. 2015, Castledine et al. 2020), and a more systematic methodology is required (Mueller and Sachs 2015, Sbahi and Di Palma 2016, Toju et al. 2020).

There are many empirical examples of multistability in community assembly dynamics (see, e.g., review by Schlöder et al. 2005). In aquatic microbial communities, Drake (1991) showed the effect of the sequence of species invasions on final community composition. More recently, Pu & Jiang (2015) found that alternative community states were maintained for many generations despite frequent dispersal of individuals among local communities. On the other hand, studies of multistability in ecology have been mainly developed for low dimensional systems (May 1977, Scheffer et al. 2001, Beisner et al. 2003). One typical example is the relationship between the abundance of phytoplankton and plant species with phosphorus concentration in lake systems (Scheffer and Jeppsen 2007); there are contrasting conditions represented by low and high algal density, and a gradual shift of phosphorus concentration triggers a rapid transition known as a catastrophic regime shift. These studies raise two important challenges regarding the analysis of compositional stability in multispecies communities, which we focus on here.

First, the potential landscape description, called a “ball and cup diagram”, has been frequently used to explain the stability of low dimensional systems (see, e.g., Scheffer et al. 2001), but how should it be interpreted when the system cannot be simplified to a few dimensions? As a solution, we introduce the concept of the *stability landscape* (Figure 1) in this paper. Walker et al. (2004) used the term to refer to the ball and cup diagram itself. Recent studies implement this idea to study the stability of microbial communities by projecting their abundance data on to a continuous potential landscape with a few dimensions (Gibson et al. 2017, Shaw et al. 2019). Here, we define the stability landscape as a structure that maps out the overall compositional stability of an ecological community. It can be represented as a graph with a set of community compositions and transition paths between them. Such a view has already been introduced in some experimental (Weatherby et al. 1998, Law et al. 2000, Warren et al. 2003) and theoretical studies (Law and Morton 1993, Capitan et al. 2011), but our goal here is to develop a methodology to study stability landscapes of empirical communities.

**Figure 1.**
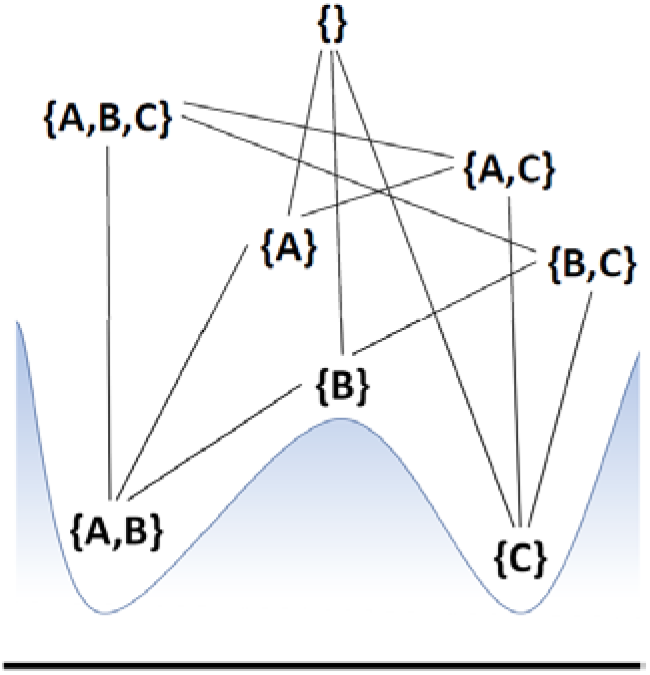
Example of a stability landscape for a conceptual three-species system. Two species, A and B, are cooperative, and these species can exclude species C if both of them are present. On the other hand, C excludes either A or B as long as they are present alone. Arrows indicates the transition of community composition when species always joins the system one-by-one. Transition from {A, C} (or {B, C}) to {A, B, C} is only valid when {B} ({A}) appears before exclusion occur. Here, both composition {A, B} and {C} is a stable state correspond to the bottom of the valley in the ball and cup diagram. The remaining six transient states can be divided into those in the basin of either stable state (in terms of the result of competition among the species present) and those on the ridge between the basins (which can transition to either of the stable states depending on the subsequent recruitment). Thus, the potential landscape represented as a smooth curve in the ball and cup diagram is embodied as a network connecting different species compositions when capturing compositional stability.

Second, it is also important to capture changes in stability landscape in response to environmental changes. For example, Lahti et al. (2014) showed that there are some taxa in human gut microbiota that show contrasting abundances according to age and lead to a shift of microbial composition during middle to old age. In this case, age was regarded as a dominant parameter of the intestinal environment and the key to transition between two contrasting states. However, given that the composition of microbiota may shift due to causes other than aging, such as infections (Kelly and LaMont 2008, Costello et al. 2012), it would be important to address changes in the overall compositional stability rather than the stability of two contrasting states.

To analyze the stability landscape of multispecies communities addressing the points described above, we introduce an *energy landscape analysis* (Becker and Karplus 1997, Wales et al. 1998, Watanabe et al. 2014a) incorporating an *extended pairwise maximum entropy model*. The extended pairwise maximum entropy model is a combination of two models that have previously been used in ecology. One is the species distribution model known as MaxEnt (Philips et al. 2004, 2006), and the other is the pairwise maximum entropy model (Schneidman et al. 2006; in ecology it is introduced as Markov network, see, e.g., Azaele et al. 2010, Araújo et al. 2011, Harris 2015). MaxEnt was developed around a model that predicts species distribution from environmental conditions, and by using the maximum entropy principle, it provides least-biased distribution estimates given prior knowledge such as observational data (Jaynes 1982, Harte and Newman 2014). The pairwise maximum entropy model has been used to predict species distribution and/or to detect signals of biotic interactions behind co-occurrence data (Azaele et al. 2010, Araújo et al. 2011, Harris 2016), and is obtained by applying the maximum entropy principle to co-occurrence information (Azaele et al. 2010).

While MaxEnt is known to give more accurate predictions relative to comparable models (Elith et al. 2006), it is unable to handle co-occurrence information that signals biotic interactions as well as shared environmental preferences between species (Kissling et al. 2012, Ockendon et al. 2014, Thuiller et al. 2015, Barner et al. 2018, Freilich et al. 2018). When community-level occurrence data is available, such information can be used to obtain more accurate predictions for species distribution (Meier et al. 2010, Leach et al. 2016, Barbaro et al. 2019). Therefore, the integration of the two models is a natural extension to handle environmental and co-occurrence data simultaneously. This has been attempted in one previous study, aimed at improving the accuracy of species distribution predictions and refining the estimation of biotic interactions under environmental heterogeneity (Clark et al. 2018), but the purpose of introducing the integrative model here is different.

By introducing the energy landscape analysis, we show how we can use the pairwise maximum entropy models to study the stability landscape of multispecies communities. While the stability landscape represents the actual compositional stability driven by population dynamics, the *energy landscape* is its approximation based on the maximum entropy principle given observational data. Briefly, energy landscape is a weighted network whose nodes represent unique community compositions and links represent transition path between them. Energy landscape analysis is the analysis of topological and connection attributes of the weighted network (Watanabe et al. 2014a). This analysis informs how a stability landscape constrains the compositional dynamics just as the ball and cup diagram does for low dimensional systems. Energy landscape analysis has its origins in the study of molecular dynamics (Becker and Karplus 1997, Wales et al. 1998) and was recently proposed as a data analysis method for neuroscience (Watanabe et al. 2014a,b, Ezaki et al. 2017, Watanabe & Rees 2017, Ezaki et al. 2018). Watanabe et al. (2014a) described the activity of multiple brain regions as binary vectors to show that the activity patterns of the resting brain are attracted to a small number of attractive states, and these states are hierarchically structured. This was done by 1) defining a set of unique activity patterns (network nodes) and connections between them as the transition path between different activity patterns (network links), 2) weighting each activity pattern by energy given by a pairwise maximum entropy model, and 3) analyzing the connection attributes of the weighted network.

We attempt to fully ground the concept of energy landscape analysis in an ecological context, and, by introducing the extended pairwise maximum entropy model, add a methodological advancement to account for the shift of stability landscapes across environmental changes. This will be a practical tool to study the compositional stability of multispecies communities in a changing environment, and will open up the possibility of empirical studies that integrate the concept of alternative stable states in community assembly studies and regime shifts developed mainly for low-dimensional systems. To show that our proposed method can be applicable to real-world ecological communities, we simulated the assembly dynamics driven by population level processes, and show that the features characterizing the stability landscape can be effectively inferred by pairwise maximum entropy models with the help of energy landscape analysis.

This paper is organized as follows. In the next section, we thoroughly explain our methodology, as well as the setup of a Lotka-Volterra (LV) model that we used for benchmarking. We then demonstrate how a stability landscape of the LV model was studied. We also benchmarked our methodology for multiple independent data sets obtained from different simulation conditions. Finally, as the application to a real-world community, we applied our methodology to the murine gut microbiota (Nakanishi et al. 2020). We found two alternative stable states and revealed how change in their relative stability is affected by age interacting with species relationships.

## Materials and Methods

### Definition of state space

By formalizing the concept of the stability landscape, we obtain a definition of the state space for an energy landscape. We define community composition σ as a binary vector of length S. Here, S is the total number of species. There is a total of 2^S^ unique community compositions, which are the nodes of the stable state. We denote a community composition of kth sample (k ∈ {0,1,…,2^S-1^}) as 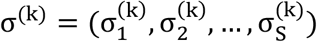, where 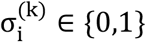 is the presence absence status of the ith species. Then, we define the links that connect community compositions under the assumption that community compositions change in a stepwise manner, i.e., two community compositions are adjacent by a link only if they have the opposite status (i.e., 0/1) for just one species. Under this assumption, the community compositions form a regular network in which each node has S links.

### Pairwise maximum entropy models and Energy landscape

We assign energy to each community composition and impose the potential structure to the state space by introducing the extended pairwise maximum entropy model. The model determines the probability of the occurrence of community composition σ^(k)^ in an environmental condition ε = (ε_1_, ε_2_,…, ε_M_), which is an array of continuous values representing environmental factors such as resource availability, pH, temperature, or age of host organism:

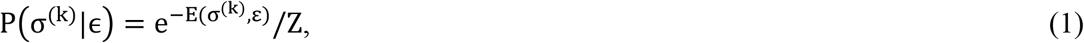

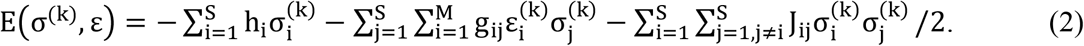

Here, E(σ^(k)^, ϵ) is defined as the energy of community composition σ^(k)^, and

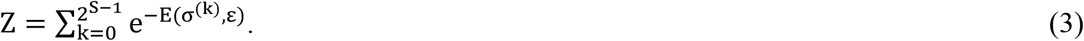

It is worth noting that we label E(σ^(k)^, ϵ) as energy simply because of the terminology for the same equation in statistical physics (Brush 1967, Azaele et al. 2010). In ecology, it is nothing but an exponent in eq.(1) and is an indicator of how often a community composition is likely to be observed; it does not correspond in any way to the physical form of energy used in the ecological studies. Parameters in eq.(2) are h_i_s, J_ij_s and g_ij_s, which are elements of a vector h = (h_1_,h_2_,..,h_S_) and matrix J = (J_ij_)_i=1,2,…S;j=1,2,…,S_ and matrix g = (g_ij_)_i=1,2,…,M;j=1,2,…,s_, respectively. h_i_ is the net effect of unobserved environmental factors that may favor the presence (h_i_ > 0) or absence (h_i_ < 0) of species i, and g_ij_ represents the effect of the ith (observed) environmental factor on the occurrence of the jth species. Each species is coupled to all others in a pairwise manner through J_ij_s (J_ij_ = J_ji_), therefore J_ij_ > 0 favors the co-occurrence of species i and j and J_ij_ < 0 disfavors the co-occurrence of species i and j. The extended pairwise maximum entropy model can be reduced to the pairwise maximum entropy model (Azaele et al. 2010, Araújo et al. 2011, Harris 2015) with,

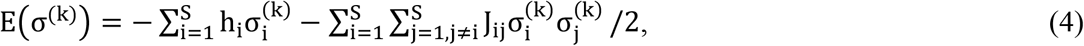

if the effect of environmental conditions can be ignored. Eq. (2) and (4) include pairwise relationships. Thus, it is assumed that all higher-order occurrence patterns are well-captured by the information encapsulated in the first two moments (Azaele et al. 2010). This is relaxed by including the high order terms in these equations. However, such extension would increase the dataset size required to obtain enough predictive performance (Nguyen et al. 2017).

The pairwise maximum entropy models assign energy to each community composition in state space. The energy represents the directionality of the transition between community compositions. For each pair of two adjacent nodes (say, σ^(k)^ and σ^(k′)^), let P(σ^(k)^) < P(σ^(k′)^) such that E(σ^(k)^) > E(σ^(k′)^). The transition of community compositions is more likely to occur from σ^(k)^ to σ^(k′)^ than σ^(k′)^ to σ^(k)^. Since such a local transition rule governs the transition dynamics in a chain, the energy landscape determines the directionality of overall compositional dynamics. We expect that this could be an approximation of the stability landscape.

We assume that we have a dataset containing the community compositions of N samples (Fig. 2A, B). We denote the community composition of samples as an S × N matrix X = (x_1_,x_2_,…,x_N_) and x_i_ ∈ {σ^(1)^,σ^(2)^,…,σ^(2S)^} (Fig. 1B). If available, we denote the environmental condition of N samples as M × N matrix Y = (y_1_,y_2_,…,y_N_), where local environmental condition 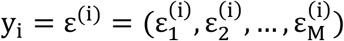 represents the environmental factors of the ith sample (Fig. 2B). Thus, a pair of X and Y is the observational data required for our analysis (Y is not always necessarily). The maximum likelihood estimates of h, J and g (Fig. 2C) can be obtained by a gradient descent for the pairwise maximum entropy model (eq.(4)) or stochastic approximation method for the extended pairwise maximum entropy model (eq.(2)) as described in Appendix S1. Since these equations are derived from the maximum entropy principle, probability distribution of community compositions with these parameters is the least biased estimate given the observational data (Jaynes 1982, Harte and Newman 2014). In other words, the probability distribution satisfies the constraints imposed by the observational data while maximizing the remaining uncertainty. This is a reasonable and well-verified (Shipley et al. 2006, Elith et al. 2006, Franklin 2009, Parisien and Moritz 2009, Harte 2011, Staniczenko et al. 2017, Clark et al 2018) assumption to estimate the probability of the occurrence of a community composition that is not actually observed.

**Figure 2.**
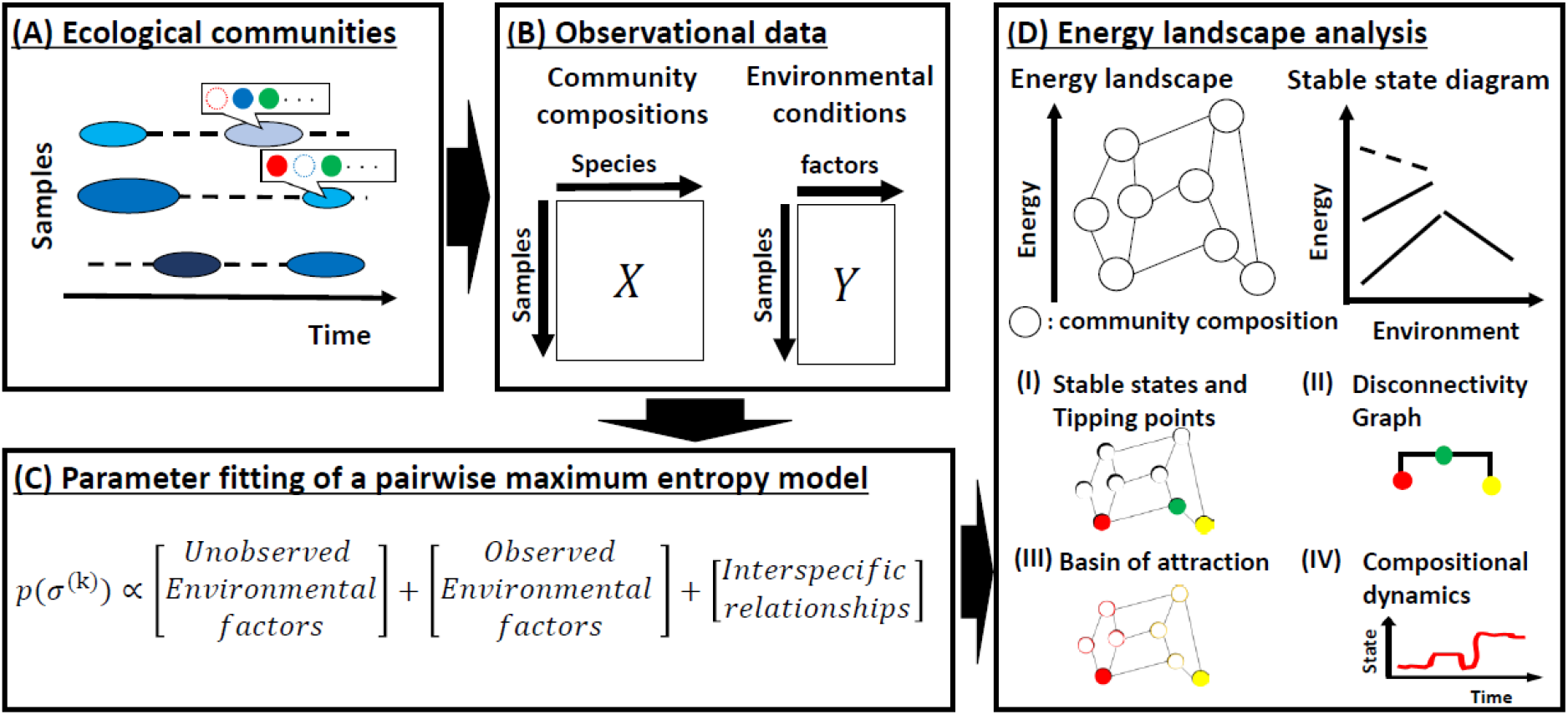
Illustrative explanation of our approach. (A) we assume that the dataset includes occurrence of species in local communities sampled from multiple targets (e.g., sites, hosts) and/or timepoints, with possibly accompanying values representing local environmental condition (environmental factors) (in the illustration, circles and filled circles show species that are absent from or present in a local community, and colors and size of ellipses represent differences in environmental condition). (B) dataset is converted to matrices of presence/absence status, and environmental factors (if they are available). (C) these matrices are used to fit parameters in a pairwise maximum entropy model. Here, P(σ^(*k*)^) is the probability of a community state σ^(*k*)^ (see Materials and Methods for the detail). (D) the fitted pairwise maximum entropy model specifies an energy landscape which is a network with nodes representing community states and links representing transitions between community compositions. Its change over environmental conditions can be described as a stable state diagram. Energy landscape analysis, acknowledges (I) the stable states (red and yellow filled-circles) and tipping points (green filled-circle), (II) disconnectivity graph summarizing the hierarchical relationships between the stable states and tipping points, (III) basin of attraction of stable states (red and yellow circles indicate basins of attraction of the two stable states), (IV) compositional dynamics constrained by the energy landscape.

### Energy landscape analysis

Energy landscape analysis is an analysis on the topological and connection attributes of an energy landscape (Fig. 2D). Below, we show how to identify its key components and explain how they can help us to understand the structure of a stability landscape. We will use the term “node” in the same sense as one community composition.

#### Energy minima

We identify energy minima of an energy landscape as *stable states* of a stability landscape (red and yellow filled circles in Fig. 2D). The local minima have the lowest energy compared to all neighboring nodes, and thus constitute end-points when assembly processes are completely deterministic (i.e., when transition of community compositions always go down the energy landscape). Presence of multistability can be identified as multiple energy minima within an energy landscape. Since an energy minimum is a node with energy less than all S neighboring nodes, we examined whether each of the 2^S^ nodes were local minima. The same idea has already been described in Azaele et al. (2010) but was there only mentioned it briefly in the analysis of a plant community distribution.

#### Basin of attraction

The basin of attraction (red and yellow circles in Fig. 2D) is the set of community compositions that reach one distinct stable state when assembly processes are completely deterministic, and are identified according to the stable state to which it converges. The basin of attraction to which a community composition belongs is determined by a steepest descent method, as follows. First, we selected a node i in the energy landscape. If the selected node is not a local minimum, we moved to the node with the lowest energy value among the nodes adjacent to the current node. We repeated moving downhill in this manner until a local minimum was reached. The initial node i belongs to the basin of the identified local minimum. We ran this procedure for each of 2^S^ nodes except for the local minima.

#### Disconnectivity graph

A disconnectivity graph is a tree diagram that summarizes the hierarchical relationships among stable states of an energy landscape (Fig. 2D). Terminal leaves of the tree represent the stable states and their vertical positions represent their energy values. Those of the branches represent the height of the *energy barrier* that separates the stable states belonging to the two branches. Here, the community composition connecting two branches is a *tipping point*. The tipping point is the lowest part of the ridge between two basins (green filled-circles in Fig. 1D). When we consider the transition from a stable state σ^(A)^ to stable state σ^(B)^ (σ^(A)^ → σ^(B)^), the height of the energy barrier is calculated as the energy of the tipping point minus that of σ^(A)^. Similarly, the height of the energy barrier for σ^(B)^ → σ^(A)^ is calculated as the energy of the tipping point minus that of σ^(B)^. Thus, the directionality of the transition between two stable states is typically asymmetrical and transitions with smaller energy barriers occur more frequently than the opposite direction. Tipping points can be obtained by checking connectivity of an energy landscape while changing an energy threshold value as follows. First, we set an energy threshold value, denoted by E_th_, to the energy value of the node that attained the second highest energy value among community compositions. Second, we removed the node (and the links connected to the node) with the highest energy value such that the node with energy E_th_ becomes the highest energy state. Third, we checked whether each pair of stable states was connected in the reduced network. Forth, we lowered E_th_ to the second highest energy value again. Then, we repeated this process until all the local minima were isolated. For each pair of stable states, we recorded a community composition having the lowest E_th_ value below which the two stable states were disconnected. In practice, some stable states must be removed because they are implausible. Here, we pruned stable states whose shallowest energy barrier was 20% lower than the highest energy barrier among all energy barriers across the energy landscape.

#### Stable state diagram

A stable state diagram unfolds the energy of stable states and tipping points in disconnectivity graphs over environmental conditions (Fig. 2D). It maps out how an energy landscape changes in response to environmental gradients. A stable state (and tipping point) is represented as a line segment that indicates a range of environmental conditions in which it is identified. Typically, we identify stable states as solid lines and tipping points as dashed lines.

#### Numerical simulations

We carried out numerical simulations to generate transition sequences of community compositions constrained by an energy landscape, and we refer to these simulations as *emulated compositional dynamics* (Fig. 2D). We employed the heat-bath method (also known as Gibbs sampling; Gilks et al. 1996) as follows. First, we selected an initial community composition. Then, at each time step, a transition from the current community composition σ^(k)^ to one of its S adjacent community composition σ(^k′)^, selected with probability 1/S, was attempted (σ^(k)^ and σ(^k′^) differ only with respect to the presence/absence status of one of S species). The transition to the selected community composition took place with probability e^−E(σ(k′))^/(e^−E(σ(k)^) + e^−E(σ(k′)^)).

By using the algorithm, we collected transition sequences of community compositions between different stable states. We then extracted the community composition having the highest energy in each transition sequence. These compositions constitute the *effective boundary* between the basin of stable states on the energy landscape. We refer to a fraction of effective boundary that mediates most of the transition between two stable states as *transition channels*.

### Competitive Lotka-Volterra model

To test the applicability of our method to community assembly dynamics driven by population level processes, we used the following Lotka-Volterra (LV) competition equation:

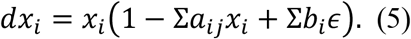

Here, *x_i_*, is the population abundance of species *i* and it is an element of abundance vector 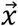. A positive real number *a_ij_* represents competitive interactions between species i and j, which is an element of an interaction matrix *A* = (*a_ij_*). The diagonal elements of *A* i.e., *a_ii_*s, represent the intraspecific interactions. The environmental condition is represented by *ϵ*. It can include multiple environmental factors, but we assumed that it is a scalar with a range of [0,1] throughout our simulation. If we do not consider environmental variation then *ϵ* = 0. Response of species *i* to *ϵ* is represented by *b_i_*, which is an element of a response vector *b*. When *ϵ* = 0, eq. (5) is considered to be one of the simplest multi-species systems whose multistability has already been investigated (Gilpin and Case 1976). In a more general view, LV equations are derived as the continuous limit of individual based models (Wilson et al. 1993) or an approximation of patch dynamic models (Keymer et al. 2000). Therefore, the LV equation-based model will be an important starting point for testing the applicability of our method to real-world ecological communities.

To simulate assembly dynamics, we introduced the processes of “extinction” and “recruitment” into our model. Extinction is a process that reduces the frequency of species if they fall below a certain threshold (*e*) to zero, and recruitment is an operation to introduce a species that is absent from the system at a propagule size (*r*). We set *e* = 10^-5^ and *r* = 10^-4^ throughout this paper. To simplify the analysis, we set a fixed interval for recruitment *τ_r_* = 300.

#### LVdata set

We generated a LV data set that characterizes the stability landscape of a LV competition model. First, we generated 20,000 empty sites (i.e., 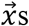 such that all elements are zero). The number of species, *S*, was fixed throughout the simulation. We assumed that the population dynamics, extinctions and recruitments occurred independently among sites. We obtained the numerical solution of the LV equation by the first-order Euler method with *dt* = 0.1. After the first species (randomly selected among *S* species) was added to each site, every *τ_r_* steps, for each site we checked the absent species, randomly select one of them and introduced it with a propagule size *r*. During these processes, the abundance vector 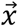 of the LV model was recorded every 10 steps after replacing the abundance of species that fell below *e* to zero. We interpreted 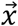 as a community composition *σ^(k)^* by an observational threshold *o* = 10^-3^, where abundance was replaced by 1 if *x_i_* > *o* else 0. Then, time series was interpreted as assembly sequences represented by community compositions. The following test for stopping the simulation was applied every *τ_r_* steps before introducing a species; let 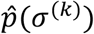 be the present probability of community compositions of 20,000 sites and 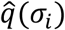 be the same value at *τ_r_* steps ago, for the set of the indices of community compositions K that appeared in records, we calculated the Jensen-Shannon divergence between them as:

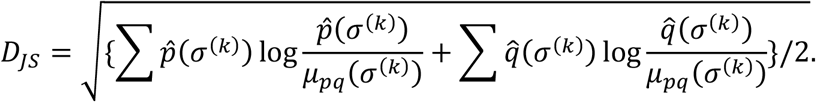

Here, 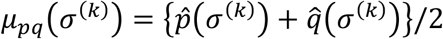. The simulation was stopped if *D_JS_* < 0.0001. We have arranged the numerical simulation procedure to facilitate the determination of stable states. It would be difficult to find an ecological interpretation of this procedure; currently, there are few studies on population-level models aimed at capturing compositional stability of multispecies communities (but see Capitan et al. 2011). It is beyond the purpose of this paper to consider what process should be implemented to reproduce the probability distribution of community compositions in nature.

#### Evaluation

By using the LV data set, we identified five features that characterize the stability landscape. Then, we compared them to the comparable features of an energy landscape inferred from community compositions sampled from the LV data set (prior to sampling, we truncated the first 25 points of the assembly sequences to remove the effect of initial conditions). For the sake of simplicity, we will refer to the former values as *actual features*. Actual features can only be obtained with large amounts of time series data. Therefore, the usual data analysis relies solely on the features of an energy landscape.

Actual features are calculated as follows. To reduce variation, we calculated these features for community compositions observed more than 20 times. Prior to this, we truncated the first five points of the assembly sequences to remove the effect of initial state. First, we identified the unique community compositions found at the end of each time series as stable states. These are directly comparable to the stable states of the energy landscape. Second, we calculated the *empirical probability* of the occurrence of a community composition σ^(k)^ across overall assembly sequences as 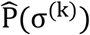. It is comparable to the probability calculated from a pairwise maximum entropy model, P(σ^(k)^), which we refer to as the expected probability. Third, we calculated the *relative convergence time (RCT)* for each community composition. This value indicates distance from a community composition to a stable state to which it converges. RCT is calculated as follows: first, if there are sub-sequences of the same composition in an assembly sequence, we removed the redundancy by replacing them with one (i.e., {…, σ^(A)^, σ^(B)^, σ^(B)^, σ^(C)^,… } → {…, σ^(A)^, σ^(B)^, σ^(C)^,…}). Then, we rescaled the position of community composition as a value between 0-1. For each community composition, RCT(σ^(k)^) is the mean of their rescaled position in all sequences in which they appeared. RCT is comparable to the energy of community states; more precisely, we compared it to the *rescaled energy* 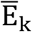, which takes 0 at a stable state and 1 at the community composition that has the highest energy within a basin of attraction. Fourth, we identified the basin of attraction of σ^(k)^ as follows: for each community composition σ^(k)^, we picked the end state of the sequences in which it appeared and counted the number of each stable state. We determined that σ^(k)^ is in the basin of attraction of a stable state to which it most frequently converged (if there is more than one such state, it belongs to all of them). It is directly comparable to the basin of attraction inferred by the energy landscape. Finally, we calculated the *imbalance score (IS)* to evaluate how stable states that a community composition converges to are uniquely determined. Similar to determining the basin of attraction, for each community composition σ^(k)^, we picked the end state of sequences in which it appeared. Then, we calculated a Gini-Simpson index for the set of stable states as the imbalance score (i.e., 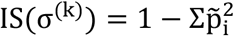; here, 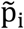 is the relative frequency of stable states). Different from other features, instead of defining a comparable value in the energy landscape, we used IS to evaluate the relevance of the hierarchical structure of the energy landscape to the actual transition dynamics.

### Murine gut microbiota

We applied our approach to the data of gut-microbiota taken from the feces of six male C57BL/6J mice, which are in the DDBJ database (http://trace.ddbj.nig.ac.jp/DRASearch/) under accession number DRA004786 (Nakanishi et al. 2020). Feces were sampled once every 4 weeks between 4 to 72 weeks of age, thus 18 data points were obtained per mouse. Hence, 108 data points are available. We transformed the relative abundance data into presence/absence data by setting a cutoff level at 1%, and we picked OTUs that were found in 20% to 80% of samples. This provided presence/absence data for 8 OTUs specified at the genus level. We also used mouse age (4-72 weeks) as an explicit environmental parameter. In the analysis, we scaled 4-72 weeks to a value within a 0-1 range. We assumed that a 4-week interval was sufficiently longer than the transient dynamics of the gut microbiota (Gerber 2014), and treated microbiota composition of the same mouse at different ages as independent data.

### Software

We used Mathematica (version 10.2 and 11.0, Wolfram Research, Inc., Champaign, Illinois, USA) to implement the method and generate simulation data. Computer codes (Mathematica notebook and package file) are available as online supporting information (DataS1).

## Results

### Analysis of a competitive Lotka-Volterra system

We analyzed the stability landscape of a 16-species competitive LV system with a fixed environmental condition (*ϵ* = 0) having three stable states (Fig 3; see Appendix S2: Table S1 for the parameter values). We first generated the LV data set, calculated actual features, and sampled 256 compositions across assembly sequences to fit the parameters of the pairwise maximum entropy model. At the population level, there were sequences of extinctions and invasions, and the system finally converged to one of three stable states. Below, in order to emphasize that we are considering community compositions, we denote a community composition as C_k_ instead of σ^(*k*)^; k is an integer obtained by transforming a binary vector into a decimal number, e.g., if the community composition is (0,1,1,0,1), it is represented as C_13_.

**Figure 3.**
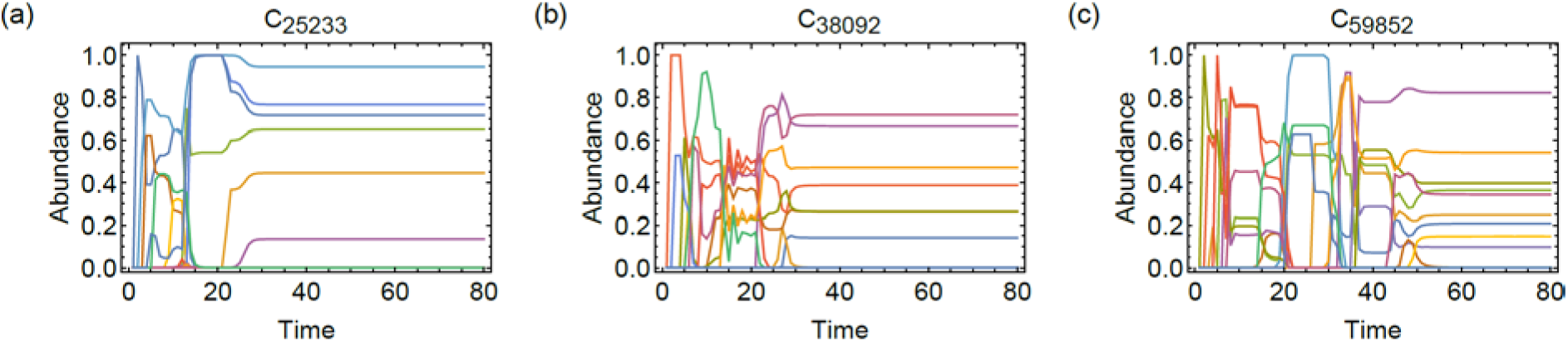
Examples of population dynamics that converged to three different stable states. (a) C_25233_, (b) C_38092_ and (c) C_59852_. See table 1 for the detail of community compositions.

The stable states were C_25233_, C_38092_ and C_59852_ and they appeared 5,332, 6,002, and 8,666 times within 20,000 simulations. We found 19,190 unique community compositions in the simulation out of 2^16^ = 65,536 possible compositions. The length of the assembly sequences was 80 points and we used 75 points after truncating the first 5 points. Thus, we had 20,000 × 75 = 150,000 data points for calculating actual features. We focused on 1628 community compositions that appeared 20 times or more. While they are only 8% of the unique community compositions found in the assembly sequences, they covered 75% of observations, *i.e*., a small fraction of community compositions appeared repeatedly.

#### Energy landscape of the LV system

Figure 4a shows the disconnectivity graph obtained from the energy landscape analysis of the LV system. Since we do not consider environmental variation in this case, we used eq. (4) for the analysis. The stable states identified by the energy landscape analysis were in perfect agreement with that of the LV system. Figure 4b is a scatterplot comparing the expected probability P(σ^(k)^) calculated by the pairwise maximum entropy model (x-axis) and the empirical probability 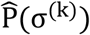 (y-axis). This analysis is not specific to the energy landscape analysis but is a general criterion to assess the performance of maximum entropy models (Azaele et al. 2010). The Spearman rank correlation coefficient was 0.40 (p<0.001). This value shows the overall performance of the maximum entropy model to predict the occurrence of community compositions. However, this value may be underestimated because the number of observations is not large enough to make an accurate calculation for 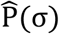 of rare community compositions. Figure 4c shows the relationship between the rescaled energy 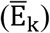 and relative convergence time (RCT; y-axis). The Spearman rank correlation coefficient was 0.36 (p<0.001), 0.69 (p<0.001), and 0.52 (p<0.001) for predicted basins of attraction of C_25233_, C_38092_ and C_59852_ and their weighted mean according to the number of observations (Table 1) was 0.55. The correlation found here indicates that the energy landscape agrees well with the actual stability landscape, in the sense that it will predict the order in which community compositions appear during transient to stable states.

**Figure 4.**
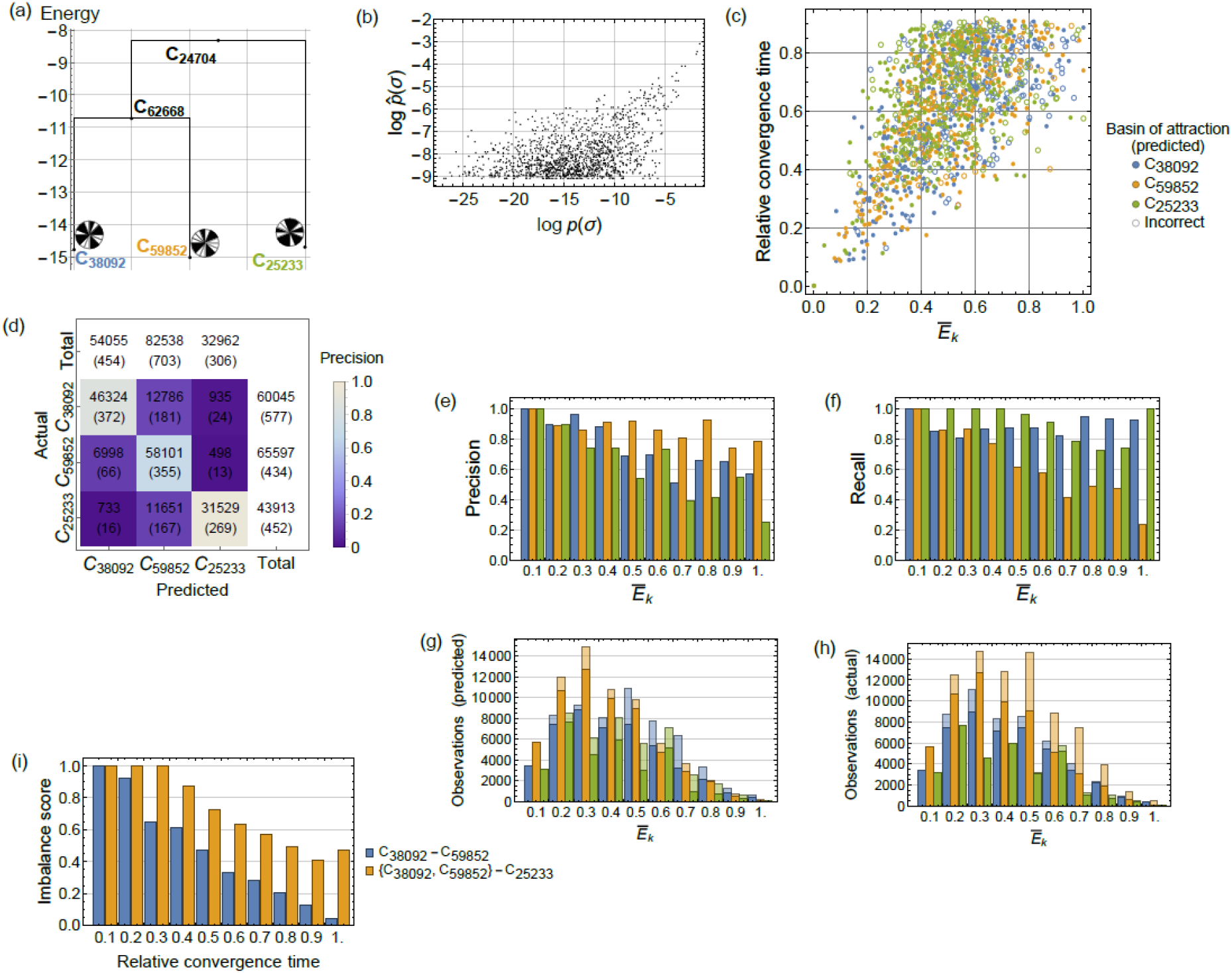
Energy landscape of a competitive LV system. (a) disconnectivity graph with three stable states and two tipping points. (b) scatterplot of the expected (P(σ^(k)^)) and empirical 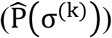 probability for community compositions. (c) scatterplot of the rescaled energy 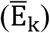 and RCT for community compositions. The color of each circles represents the basin of attraction of the energy landscape and filled or open circles indicate whether it agrees (or disagrees) with that of the stability landscape. (d) confusion matrix with the number of observations (the number of community composition is shown in parentheses). Here, observations of stable states were removed from the result. (e) precision of the prediction of basin of attraction with respect to observations. (f) recall of the prediction on basin of attraction with respect to observations. (g) total number of observations for predicted basin of attraction. (h) total number of observations for actual basin of attraction. In (e-h), values were calculated for 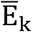 value grouped into 10 bins of equal width. Color of bars indicate predicted basins of attraction in (g), and actual basins of attraction in (h). Shaded and unshaded areas in (g, h) indicate observations with mismatch and match between actual and predicted basins of attraction, respectively. Shaded areas are stacked on the unshaded areas. The ratio of the unshaded areas to the total length of the bars corresponds to precision in (g) and the same value in (h) corresponds to recall. (i) imbalance score calculated for the balance between C_38092_ and C_59852_ and C_25233_ and C_38092_-C_59852_. In the latter case, C_59852_ identified with C_38092_.

**Table 1.** Predictability of stable states.

| Stable states | Composition | Number of observations | Basin of attraction (precision) | Basin of attraction (recall) |
| --- | --- | --- | --- | --- |
| C<sub>25233</sub> | (0,1,1,0,0,0,1,0,1,0,0,1,0,0,0,1) | 5332 | 0.72 | 0.96 |
| C<sub>38092</sub> | (1,0,0,1,0,1,0,0,1,1,0,0,1,1,0,0) | 6002 | 0.78 | 0.86 |
| C<sub>59852</sub> | (1,1,1,0,1,0,0,1,1,1,0,0,1,1,0,0) | 8666 | 0.89 | 0.70 |

We next evaluated how well the energy landscape predicted the basin of attraction. The color of each circle in figure 4c represents the basin of attraction of the energy landscape. A filled or open circle represents whether it agrees (or disagrees) with the stability landscape. The overall result for the prediction is summarized as a confusion matrix with both number of community compositions and observations (Fig. 4d). As suggested from the disconnectivity graph (Fig. 4a), the basins of attraction of C_38092_ and C_59852_ are more easily mislabled with each other than those of C_25233_ and C_38092_ or C_25233_ and C_59852_. To evaluate the predictive performance in more detail, we calculated precision and recall for different positions in the energy landscape according to the number of correct/incorrect observations (Fig. 4e-h). The overall precision and recall for the basin of attraction of C_25233_, C_38092_ and C_59852_ is in Table 1. The weighted mean of the precision and recall according to the number of observations of each stable state was 0.80 and 0.79, respectively. As the overall trend, both precision and recall increased when 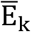 decreased (Fig. 4e,f). Since community compositions having lower 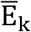 are expected to be close to the stable states, this result suggests that the energy landscape is a good approximation of the stability landscape especially around the stable states. Furthermore, in addition to figure 4d, we see how the hierarchical relationship among the stable states (fig. 4a) reflect the relationship of stable states in the stability landscape. As energy decreased, the recall for the basin of attraction of C_25233_ increases faster than that of C_38092_ and C_59852_ (fig. 4f). Moreover, the imbalance score (IS) for C_25233_ and (C_38092_, C_59852_) (here, C_59852_ is identified with C_38092_) was always larger than that of C_38092_ and C_59852_ (fig. 4i). This suggest that the transition towards C_25233_ tends to be determined at an earlier stage of assembly processes compared C_38092_ and C_59852_. In other words, these two community compositions are in close proximity in the stability landscape. These results show the agreement between the dysconnectivity graph (fig. 4a) and the numerical simulations.

#### Emulating community assembly dynamics

We considered the transition between stable states in the energy landscape to see how they inform the ridge structure that lies between the basins of stability landscape. Since we are considering assembly dynamics starting from an empty state, such ridge states may correspond to a set of community compositions where the terminal stable state is almost unique below it (i.e., they may be representing the points of no return in assembly processes). If a community composition is located on a ridge, it will have an intermediate RCT value in the sense that it is apart from both the stable states (where RCT is 0) and the states at the earlier stage of assembly dynamics (where RCT is close to 1). Also, because the dynamics are less deterministic on the ridge, it’s IS will be lower than that of other community compositions having a comparable energy.

Here, to demonstrate how we can use the energy landscape to study the ridge structure, we investigated the transitions between the stable state C_38092_ and C_59852_ in more detail. By using the heat bath algorithm, we collected 5,000 transition sequences of community compositions from C_38092_ to C_59852_ (C_59852_ to C_3809_). We then extracted the community composition having the highest energy in each transition sequence as the effective boundary between them. There were 1,616 such compositions and, as shown in Figure 5a, in the LV data set, 81% of 14,668 sequences that converged to C_38092_ or C_59852_ (Table 1) contained one of these compositions. We focus on the 100 lowest energy compositions as transition channels, since 0.3% of them (that is 56 out of 19,190, of all observed community compositions) actually appeared in 52% (7,596 out of 14,668) of assembly sequences. Figure 5b is the scatterplot showing the energy and RCT of community compositions where transition channels are highlighted as red points, and Figure 5c is the smoothed histogram of the RCT of transition channels and that of all community compositions. RCT of transition channels had a median of 0.33 (95% CI, 0.31-0.37), which was lower than that of all compositions, which was 0.65 (95% CI, 0.64-0.66). Furthermore, the IQR (difference between the third and first quartiles) of transition channels was 0.16 (95% CI, 0.10-0.22), which was lower than that of all compositions (0.29; 95% CI, 0.27-0.30). Thus, transition channels tended to be clustered in a narrow area of the lower part of the stability landscape. Figure 5d shows RCT and IS of transition channels (red) or other 130 community compositions in the range of energy where channels are distributed (Fig. 5b). The median of IS was 0.60 (95% CI, 0.56-0.79) for transition channels and it was smaller than that of others which was 0.80 (95% CI, 0.65-0.92). From the above, it can be said that there is a certain relationship between the ridge that controls the transitions between C_38092_ and C_59852_ on the energy landscape and the structure of the stability landscape that determines direction of assembly towards C_38092_ or C_59852_.

**Figure 5.**
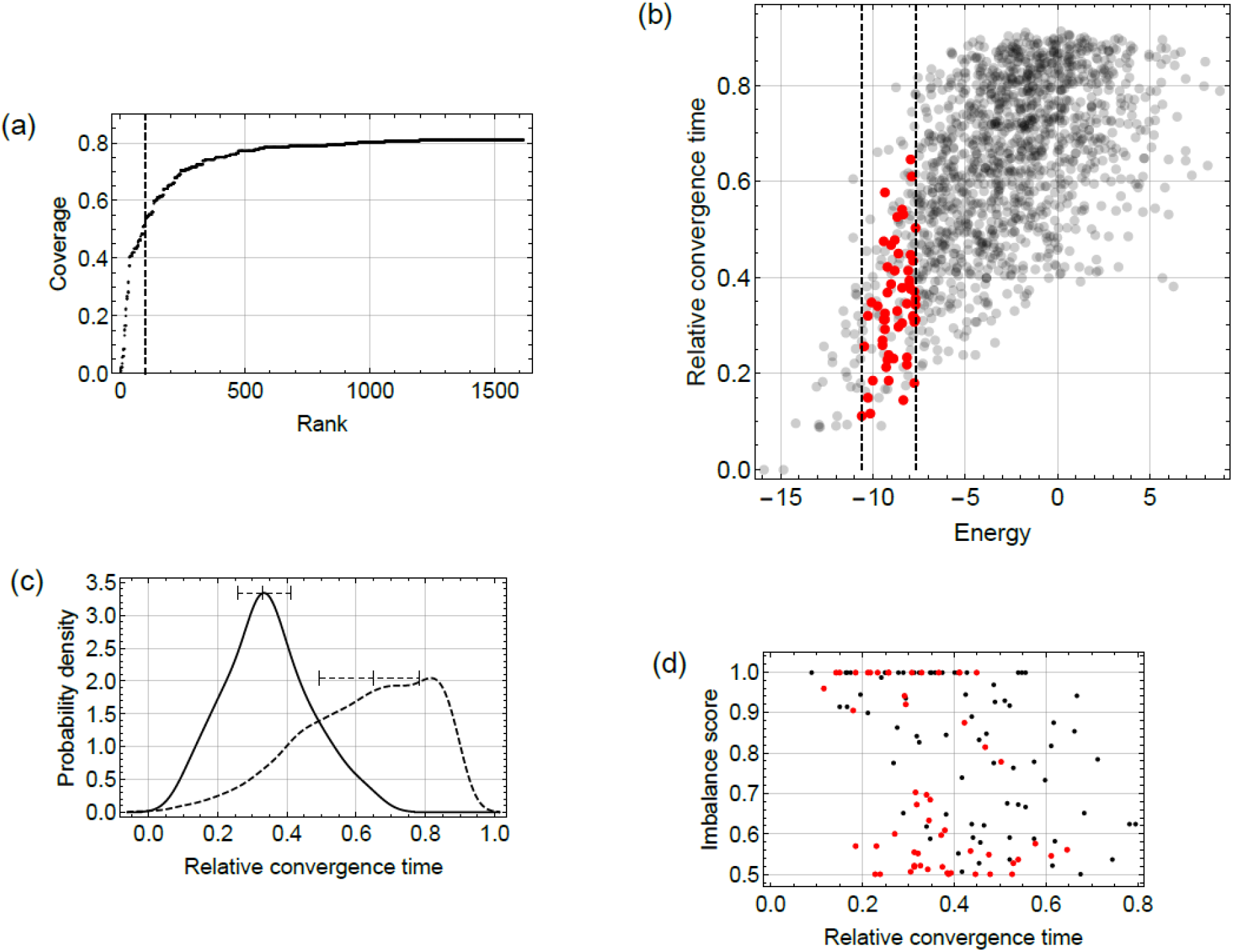
Emulated compositional dynamics between stable state C_38092_ and C_59852_. (a) coverage of assembly sequences of the LV data set that converged to C_38092_ or C_59852_ (14,668 in total) including the community compositions in effective boundary ranked by their energy. Dashed line indicates top 100 community compositions (transition channels). (b) smoothed histogram of RCT of transition channels and that of all community compositions. (c) scatterplot with energy and RCT for community compositions belonging to the basin of attraction of C_38092_ and C_59852_, where transition channels are indicated by red points. (d) relationship between RCT and IS, where transition channels are indicated by red points.

#### Energy landscape across environmental gradient

If species occurrence depends on abiotic factors, change in the stability landscape in response to environmental changes will be observed. Here, we generated LV data sets with variation in environmental parameter *ϵ* and a fixed response vector *b* (see Appendix S2: Table S1 for the parameter values), and demonstrate how the energy landscape of the extended pairwise maximum entropy model can capture the change in the stability landscape across *ϵ*.

We generated a LV data set for *ϵ* at every 0.01 steps between 0 and 1 to calculate the features of a stability landscape. Figure 6a shows the number and composition of stable states with respect to *ϵ*. The black lines show the range of *ϵ* in which a community composition was observed as a stable state. There were three branches that originated from C_59852_, C_25233_ and C_38092_; C_59852_ disappeared at *ε* = 0.08, C_25233_ changed to C_58001_ at *ϵ* = 0.125 and then disappeared at *ε* = 0.3, and C38092 sequentially changed to five other community compositions across *ϵ* = 0 to 1.

**Figure 6.**
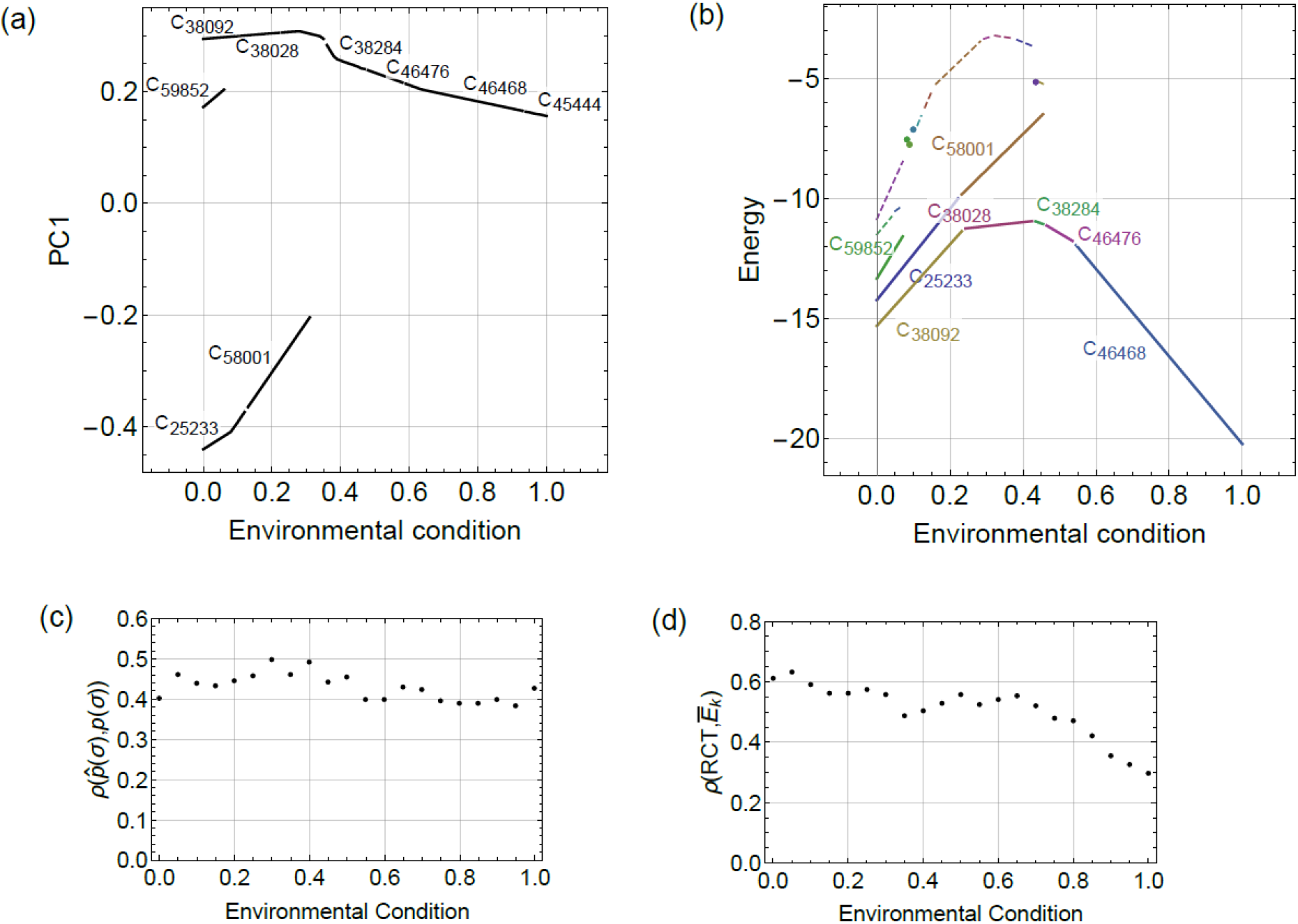
Change of stability landscape and energy landscape in response to environmental change. (a) stable state diagram calculated from LV data sets. Here, vertical axis is PC1 calculated from PCA including all stable states. (b) stable state diagram obtained from the energy landscape analysis. Solid lines indicate stable states and dashed lines indicates tipping points. (c) Spearman rank correlation between the expected (P(σ^(k)^)) and empirical 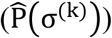 probability for community compositions. (d) Spearman rank correlation between the rescaled energy 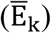 and RCT for community compositions.

We randomly sampled 256 community compositions from the above data set for the parameter fitting of the extended pairwise maximum entropy model and then applied the energy landscape analysis. Our approach identified all existing branches of stable states and slightly over-estimated the range of *ϵ* in which they exist (Fig. 6b). Also, it captured the sequence of the transition of community compositions within the branch of C_25233_ and C_38092_, except for false detection of C_41617_ and non-detection of C_38284_. The correlation between 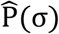 and P(σ) (Fig. 6c) was around 0.4 and slightly decreased with *ϵ*. This value was comparable to the corresponding value in the constant environment we described above (0.40). Correlation between RCT and 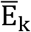 (Fig. 6d) was mostly larger than 0.5 before *ϵ* < 0.7, and then it started to decrease. While we did not investigate the reason in detail, this might be due to the increase in the stability of a single stable state that can reduce the range of community compositions other than the stable state within the data set. Overall, however, the addition of an environmental axis did not significantly reduce the accuracy of the stability landscape estimates.

### Benchmarking

In our baseline condition (A in Fig. 7), the median Spearman rank correlation between the empirical and expected probability for the community compositions was 0.43, for RCT and 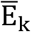 the same value was 0.51. The median of both precision and recall was 1 for stable states, and 0.79 for the basins of attraction. Reducing the number of stable states in the LV system only modestly affected the results (B in Fig. 7), but increasing it slightly made it difficult to predict basins of attraction (C in Fig. 7). This is unsurprising given that it increases the likelihood of confusion among basins. Increasing the number of species tended to make identification of stable states and their basins more difficult (D in Fig. 7). However, the difference would be small, given that the total number of possible compositions of 24-species system is 256 times larger than for the 16-species model. Although the increase of the data set size improved the results when comparing 128 and 512 data point cases (E and F in Fig. 7), no additional improvement was found when comparing 256 and 512 data point cases (A and F in Fig. 7). Because of the fundamental difference between the stability landscape of LV system and the energy landscape, this suggests the limitations of approximating the former by the later. Both Type II functional response (G) and system noise (H) did not alter the results much, but slightly reduced the correlation between 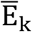 and RCT.

**Figure 7.**
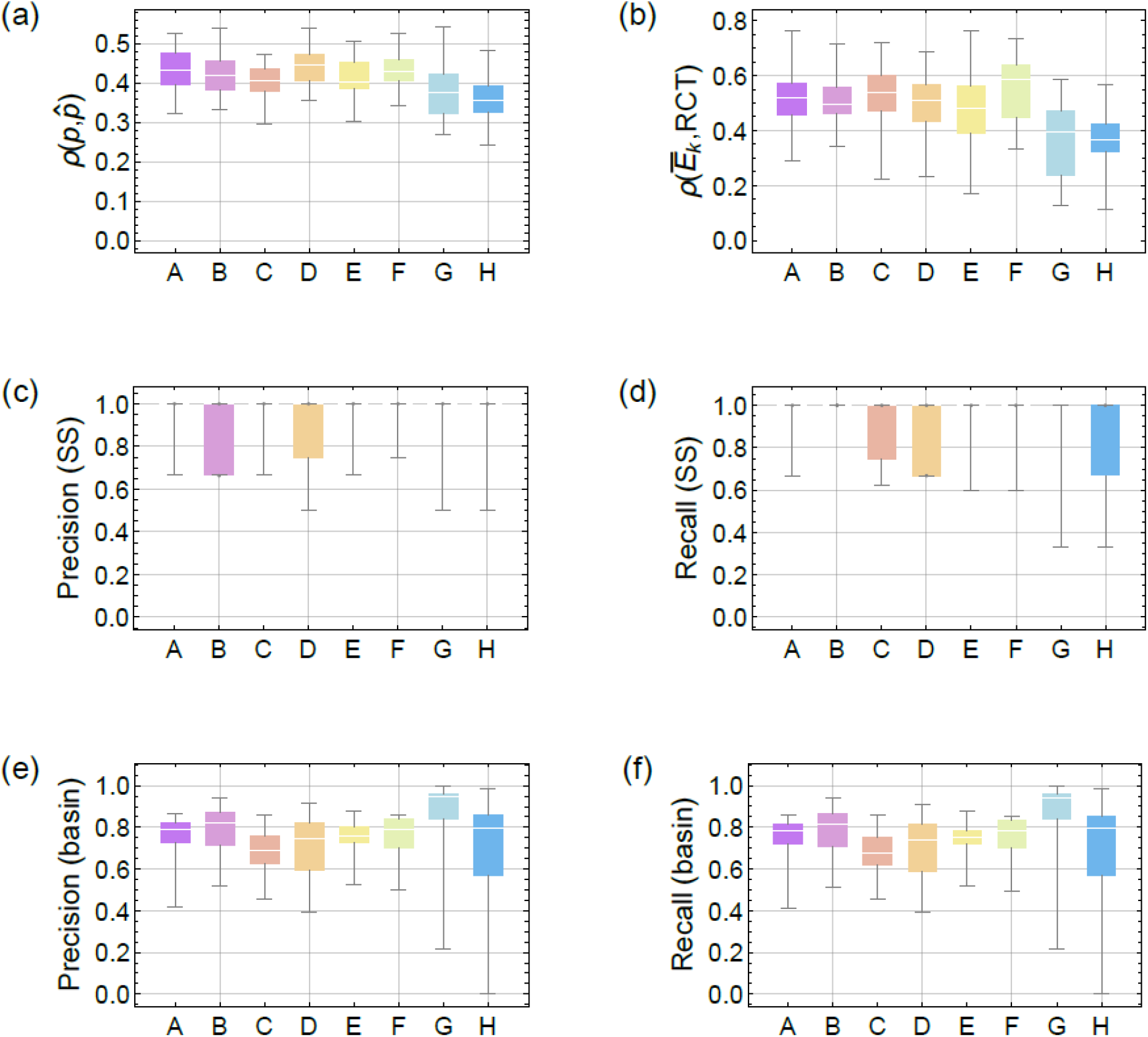
Benchmarking with different simulation conditions. Spearman rank correlation between (a) expected and empirical probability for community compositions, and (b) rescaled energy and RCT, (c, d) precision and recall for the prediction on stable states and (e, f) the same values for basin of attraction. Precision (recall) for stable states is calculated by the ratio of correctly identified states in the predicted (actual) stable states, and the same values for basin of attraction are weighted by the number of observations (see main text). The median and the first and third quartile value was 1 in A,C,E,F, G, H in (c) and A,B,E,F, C in (d). In B, D in (c) and C, D, H in (d) the median and the third quartile value was 1. We used 30 independent datasets for each simulation condition. Summarizing the simulation conditions as (number of species, number of stable states in LV system, number of samples, type of functional response, strength of noise), they are, A: (16, 3, 256, 1, 0), B: (16, 2, 256, 1, 0), C: (16, 4, 256, 1, 0), D: (24, 3, 256, 1, 0), E: (16, 3, 128, 1, 0), F: (16, 3, 512, 1, 0), G: (16, 3, 256, 2, 0), H: (16, 3, 256, 1, 0.1). Here, for the type of functional response, 1 indicates Type I and 2 indicates type II. See Appendix S2 for more information.

### Application to real data

We applied our approach to the gut microbiome data of six mice with 108 samples. Community compositions were represented by eight genus-level OTUs that can be identified as species in the above analysis, and we used mouse age as the environmental condition (i.e., only one environmental factor was considered). Since we included only age as the environmental factor in the analysis, in the extended pairwise maximum entropy model (eq. (2)), *ε* is a scalar associated with each sample and g is a vector that represents response of each species to age.

At the community level, there were two stable states C_227_ and C_93_ at initial age (Fig. 6a). C_227_ remained until 72 weeks and showed reduced energy with increasing age. C_93_ had lower energy than C_227_, and increased energy as age increased. C_93_ changed to C_125_ at 29 weeks of age and then showed reduced energy with increasing age. C_125_ further changed to C_253_ at 38 weeks of age. The energy of C_125_ and C_253_ was lower than C_227_ up to 72 weeks of age. The difference in the height of the energy barriers between the two alternative stable sates (i.e., distance between the stable states to the tipping point with respect to y-axis in Fig. 8a) decreased with increasing age until C_125_ changed to C_253_. These results suggest that C_93_ is more representative of gut microbiota during early life stages. The difference between C_93_ and C_227_ was the presence of a group of three genera (unclassified Lachnospiraceae, unclassified Ruminococcaceae and *Oscillospira*) or *Sutterella* (Table 2). Although C_93_ did not contain *Turicibacter* and *Bifidobacterium*, these genera sequentially appeared as C_93_ changed to C_253_: *Turicibacter* appeared when C_93_ changed to C_125_ and *Bifidobacterium* appeared when C_125_ changed to C_253_.

**Figure 8.**
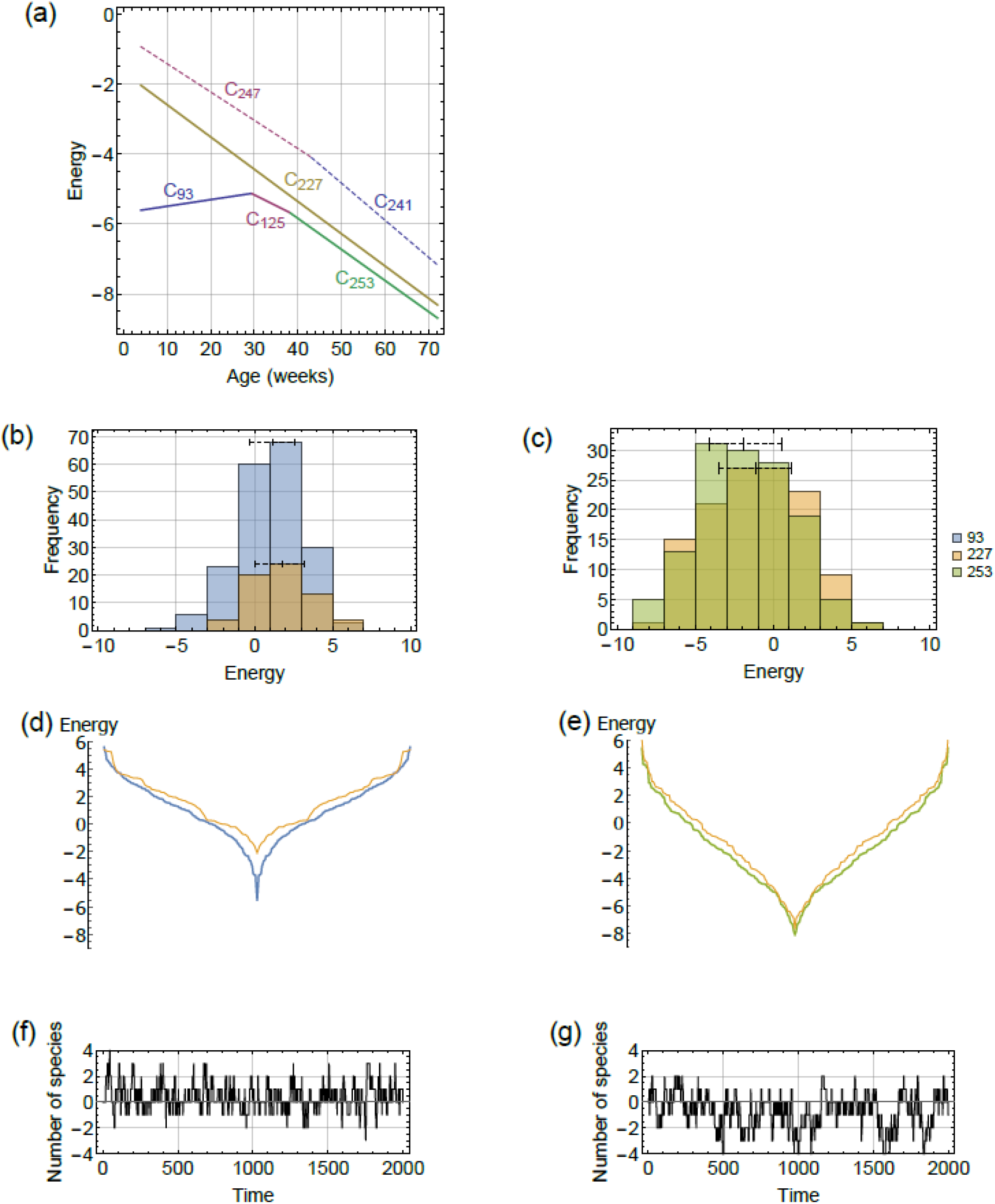
Stability of a murine gut microbiota. (a) stable state diagram showing the energy of stable states and tipping points. Here, both stable states (solid lines) and tipping points (dashed lines) are shown. Each line segment labeled by community composition represents the range of age in which stable states (or tipping points) exist. (b) the energy distribution of community compositions in the basins of attraction of C_93_ and C_227_ at 10 weeks of age. (c) the energy distribution of community compositions in the basins of attraction of C_227_ and C_253_ at 60 weeks of age. In (b, c) horizontal bars indicate the median and IQR. (d) slope of the attractive basin of C_93_ (blue) and C_227_ (yellow) in 10 weeks of age. (e) slope of the basin of attraction of C_227_ (green) and C_253_ (yellow) in 60 weeks of age. (f) emulated compositional dynamics around C_253_. (g) emulated compositional dynamics around C_253_. In (f, g) vertical axis shows the number of species of corresponding time point minus that of the stable state.

**Table 2.** Summary statistics of the energy distribution of basins of attraction.

|  | 10<br>weeks |  |  |  |  |  | 60<br>weeks |  |  |  |  |  |
| --- | --- | --- | --- | --- | --- | --- | --- | --- | --- | --- | --- | --- |
|  | Number of<br>communit<br>y<br>compositi<br>ons | Media<br>n | Min<br>(stable<br>state) | Median-Min | IQR | Kurtos<br>is | Number of<br>communit<br>y<br>compositi<br>ons | Media<br>n | Min<br>(stable<br>state) | Media<br>n-Min | IQR | Kurtos<br>is |
| C<sub>93</sub> | 191 | 1.16 | -5.59 | 6.76 | 2.90 | 2.90 | 0 | n/a | n/a | n/a | n/a | n/a |
| C<sub>227</sub> | 65 | 1.77 | -2.10 | 3.87 | 3.12 | 2.28 | 124 | -1.11 | -7.68 | 6.58 | 4.62 | 2.29 |
| C<sub>253</sub> | 0 | n/a | n/a | n/a | n/a | n/a | 132 | -1.87 | -8.08 | 6.21 | 4.66 | 2.34 |

In addition to the height of energy barriers, the number of community compositions and energy distribution of the basins of attraction provides information on the robustness and variability of stable states. Figure 8b shows the energy distribution of community compositions in the basins of attraction of C_93_ and C_227_ at 10 weeks of age. The median and IQR of the two energy distributions were almost the same, but the difference in the number of community compositions was large (Fig. 8b, Table 2). In other words, C_93_ is more stable to stochastic variation than C_227_ in the sense that it has a larger basin of attraction. However, this difference almost disappeared by 60 weeks of age (Fig. 8c, Table 2). On the other hand, the difference between the median and stable state energies was 6.76 for C_93_ and 3.87 for C_227_ in 10 weeks of age (Table 2). This was mainly because of the fact that C_93_ had a sharp decline in energy around the stable state (Fig. 8d), which is also indicated by the smaller kurtosis for C_93_ than C_227_ (Table 2). Interestingly, in C_253_, which is on a branch of C_93_ (Fig. 8a), the difference between the median and stable state energies decreased (accompanying the increase in kurtosis) (Fig. 8c, Table 2) despite the decrease in stable state energy. This resulted in a reduced slope around the stable state (Fig. 8e), and lead to increase in the amplitude and autocorrelation of compositional variation around C_253_ (Fig. 8f, g).

This is not specific to energy landscape analysis, but analysis of the estimated parameters provide some insights into the mechanisms behind community level responses (Azaele et al. 2010, Harris 2016). In figure 9a, the community level response is shown by the bars marked as ‘Total’ in addition to the genus level responses. The net effect of interspecific relationships 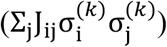, the bacterial responses to age (g_i_ϵ), and the implicit effect of environmental factors (h_i_; here, these values represent the sum of the effects other than the bacterial response to age or the biotic interaction between bacteria) are indicated by blue, orange and green, respectively. There were positive relationships among Lachnospiraceae, Ruminococcaceae and *Oscillospira* (Fig. 9b) and these relationships made up most of the interspecific relationships in C_93_, C_125_ and C_253_ (Fig. 9a). The three genera had a negative relationship with *Suterella* (Fig. 9b). Thus, they could be mutually exclusive, which was responsible for the presence of two alternative stable states over age. *Sutterella* had a positive relationship with *Turicibacter* and *Bifidobacterium*, supporting their presence in C_227_ (Fig. 9). In contrast, *Turicibacter* and *Bifidobacterium* had a negative relationship with Lachnospiraceae, Ruminococcaceae and *Oscillospira*. Thus, they were absent from C_93_ and then appeared as an effect of age. The correlation between age and the presence of a genus became positive if g (genus level response to age; Table 3) was positive; this correlation became negative when g was negative. The sum of g across community members determined the community level response to age (Fig. 8a). Since both *Turicibacter* and *Bifidobacterium* were positively affected by age, they appeared with increasing age despite their negative relationship with Lachnospiraceae, Ruminococcaceae and *Oscillospira*. In Fig. 9a the effect of interspecific relationships with *Turicibacter* in C_125_ was negative, while it was positive in C_253_ because of a positive relationship with *Bifidobacterium* (Fig 9b). The relationship between *Turicibacter* and *Bifidobacterium* also facilitated the appearance of *Bifidobacterium* at 38 weeks of age (Fig. 8a). Because of their positive relationship with age, the appearance of *Turicibacter* and *Bifidobacterium* changed the community level response to age when C_93_ changed to C_125_ and C_125_ changed to C_253_.

**Figure 9.**
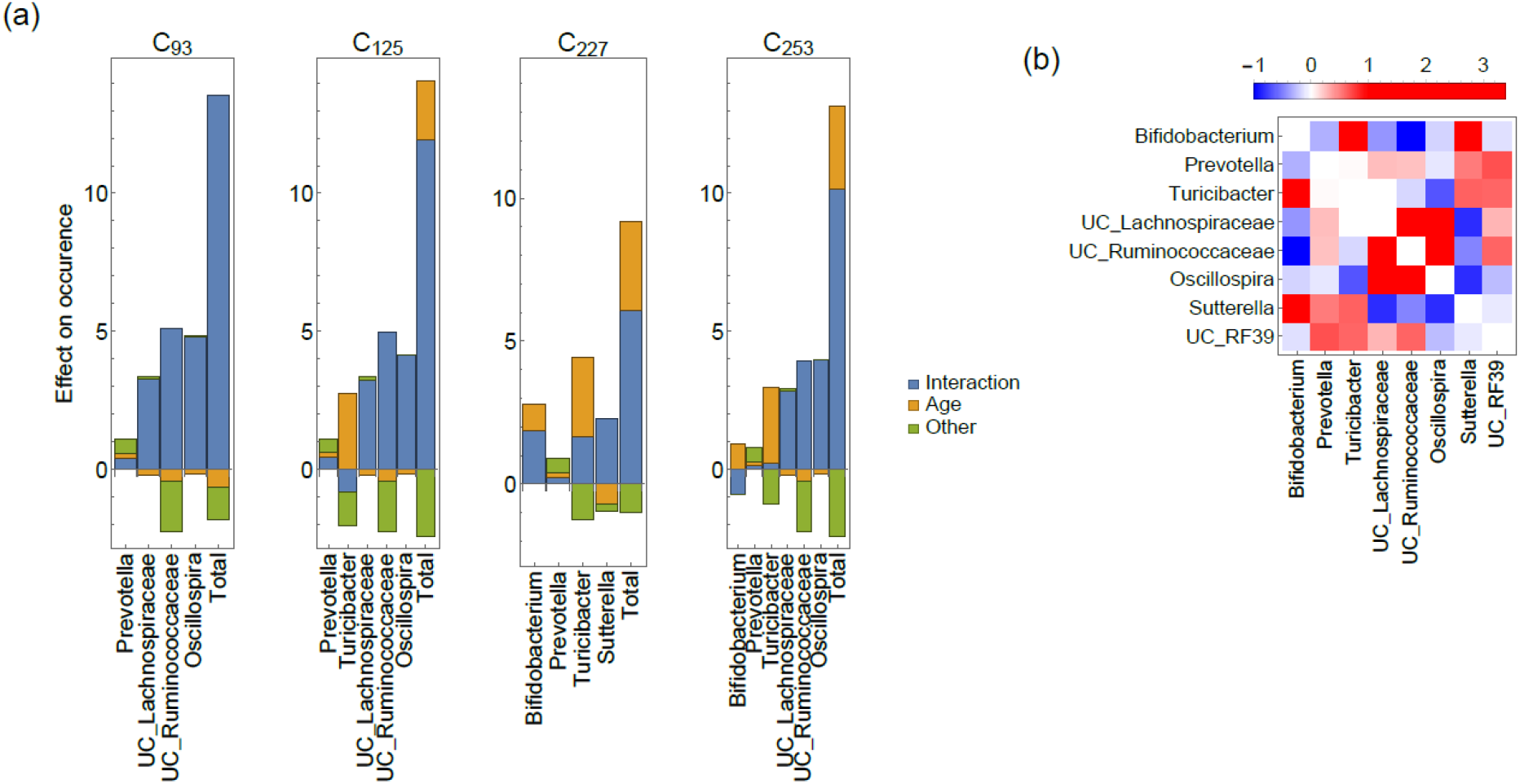
Estimated parameters for a murine gut microbiota. (a) Strength of the net effect of interspecific relationship 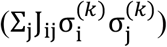, observed environmental factor (response to age) (g_i_ε) and unobserved environmental factors (h_i_) for each genus i (genus level effects) and their sum over community members (community level effects), 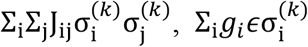 and 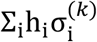 (shown as ‘total’). 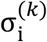 represents membership of each community (Table 3). For comparison, we set ϵ = 0.5. (b) elements of interspecific relationships (J_ij_). The value represents the strength of association between two genera (shown in columns and rows). There is a positive association between two genera if the value is positive whereas there is negative association if it is negative.

**Table 3.**
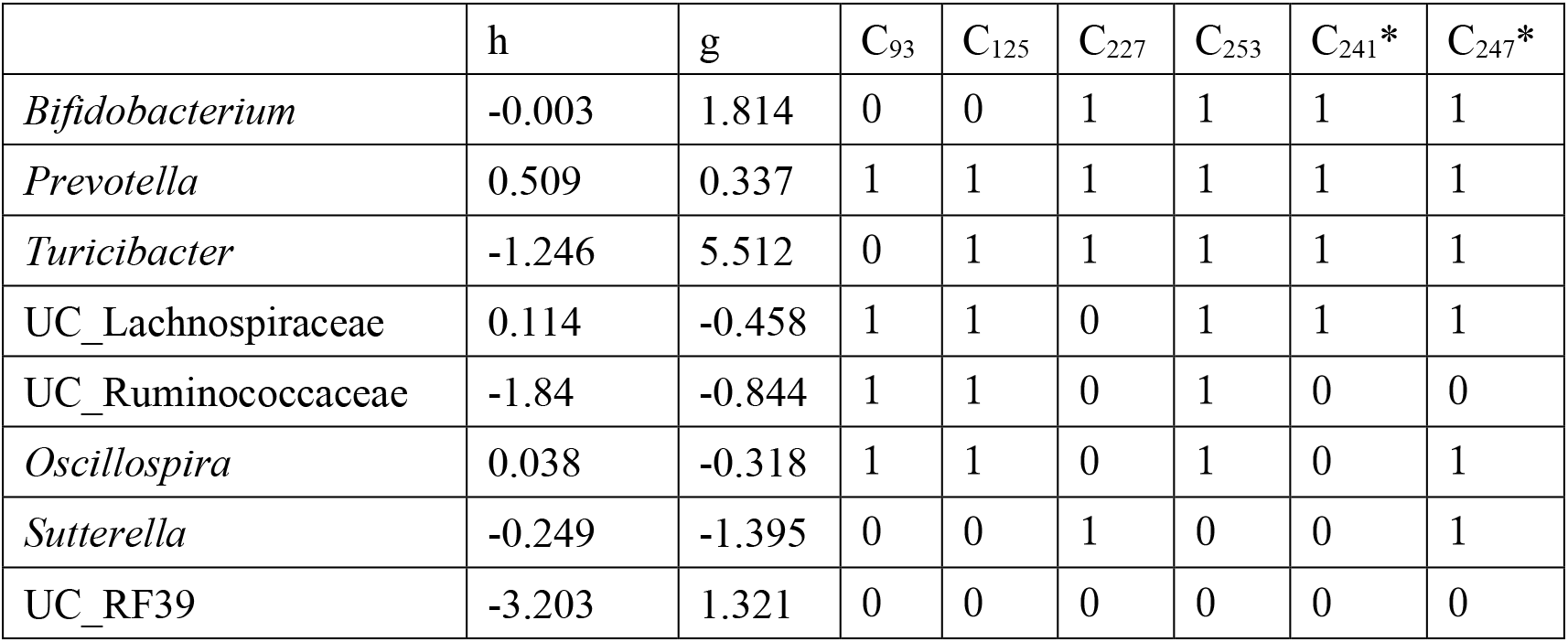
Profile of eight genera included in the analysis. h (effect from unobserved environmental factors) and g (genus level response to age) are the parameters inferred by stochastic approximation, and 0/1 values indicate membership of each genus in stable states and tipping points (marked by *).

Our results show that there were two alternative stable states and their relative stability (difference in the height of the energy barriers between them) changed over age. To examine how this result explains microbial population dynamics represented by relative abundance data, we applied analysis of similarity (ANOSIM) to abundance of the 8 genera used for the energy landscape analysis. ANOSIM statistics of the abundance data grouped with age (4-28 or 48-72 weeks) across different individuals was −884.8 (p<0.001), suggesting a difference in gut microbiota among early and later life stages. We then grouped abundance data with individuals in 4-28 or 48-72 weeks and applied the same analysis. ANOSIM statistics were - 65.2 (p<0.001) for 4-28 weeks and −37.6 (p=0.097) for 48-72 weeks. Thus, individual dependency of microbiota was strong in the early life stage and weakened in the later life stage. This was in correspondence with the structure of the energy landscape at 4-28 weeks; since the energy barrier was higher and basin of attraction is larger for C_93_ than C_227_ (Fig. 8a, b), community composition would more likely remain in the basin of C_93_ once it shifted from that of C_227_. At 48-72 weeks, switching between C_227_ and C_253_ is more likely because of the increased autocorrelation and amplitude of compositional variation (Fig. 8g) as well as the small difference in energy barrier between the two stable states (Fig. 8a), and it would reduce the individual dependency of stable states.

## Discussion

We showed that with a relatively small amount of data, our method is able to identify the features characterizing the stability landscape, i.e., it successfully detects stable states (Table. 1) and predicts basin of attraction (Fig. 4e-h), the position of each community composition with respect to the stable states (Fig.4c), and whether a community composition is on a ridge of basins in which end state is not uniquely determined (Fig.4i, Fig5). We confirmed with multiple different data sets that these results were mostly robust to different simulation conditions (Fig. 7), although the results suggested the limitation of approximating the stability landscape as the energy landscape. We also demonstrated that the same analysis was applicable to the case in which environmental conditions for each sample were different (Fig.6), without the need for increasing the amount of data. This result highlights that our method reliably captures the change of the overall compositional stability of a multispecies community over environmental change, which we regarded as the requisites for integration of compositional stability and regime shift concepts.

Our approach to quantifying compositional change in multi-stable communities could help the development of early warning signals for shifts of stable states (Scheffer et al. 2012, 2015). Our analysis suggested that a small fraction of community compositions may be channels for transitions between alternative stable states (Fig. 5). If a system is in one of such states, it may indicate an increasing probability of transition to another stable states. This would be a basis for developing an early warning signal and considering some possible intervention to prevent such shifts. Secondly, given how spatial scale interacts with the drivers of community assembly (e.g. Chase 2014), the results of energy landscape analysis should be interpreted in light of the spatial scale of total sampling extent to disentangle the drivers of community assembly across scales (Viana & Chase 2019, Ross et al. submitted). If the spatial extent of observations is large and the effect of environmental heterogeneity is significant, by choosing a relevant environmental factor (or appropriately reducing multiple factors into a single dimension), the shift of stable community compositions can be unfolded with respect to the parameter as in the stable state diagram (Fig. 6b). One could then explore whether the model’s explanation is consistent with our process-based understanding of community assembly dynamics. Such an approach would complement existing methods proposed to assess the relative importance of the ecological processes that drive community assembly (Legendre et al. 2009, Meynard et al. 2013, D’Amen et al. 2018, Mertes & Jetz 2018).

In the murine gut microbiota, our approach showed the presence of two alternative states over age (Fig. 8a). The difference between the two stable states was characterized by presence of *Suterella* or unclassified Lachnospiraceae, unclassified Ruminococcaceae and *Oscillospira* (Table 3). Our result suggested that the stable state containing the three genera (C_93_) was more representative of the gut microbiota during early life stages because it had lower energy than the counterpart (C_227_) and became relatively unstable with increasing age (Fig. 8f, g). Langille et al (2014) reported that Lachnospiraceae, Ruminococcaceae and Oscillospiraceae are phylogenetically closely related and characterize the murine gut microbiota during early life stages. Our result supports tight associations among these groups and their role characterizing a stable state representing early-life gut microbiota. C_93_ changed to C_125_ (29 weeks age) and C_125_ changed to C_253_ (38 weeks age) due to appearance of *Turisibacter* and *Bifidobacterium*, respectively. The response of *Turicibacter* and *Bifidobacterium* to age had a prominent role in the community level response during these shifts. However, the importance of interspecific relationships was also identified. Since *Turicibacter* and *Bifidobacterium* had a negative relationship with Lachnospiraceae, Ruminococcaceae and *Oscillospira*, they could not appear until 28 and 39 weeks of age, respectively. Here, presence of *Turicibacter* facilitated appearance of *Bifidobacterium* due to their positive relationship. In terms of the height of energy barriers, the transition between alternative stable states tended to be unidirectional (from C_227_ to C_93_) in the early life stage and bidirectional in the later life stage (between C_227_ and C_125_ or C_253_). Furthermore, the mechanism of changing compositional variability was inferred from the analysis of energy distribution of basins of attraction. Our results suggest that there were two mechanisms. The first is the change in the number of community compositions in the basin of attraction, which directly affects the magnitude of compositional variability a basin accommodates, and thus corresponds to Holling’s ecological resilience (Holling 1996). The second is the change in the shape of basin of attraction, especially related to the kind of curvature around the stable state (Fig. 8d, e). This relates to the return time to a stable state, and corresponds to Holling’s engineering resilience (Holling 1996). Not only the change in the height of barrier between stable states but also these mechanisms governing compositional variability works synergistically in the age-related loss of stability in murine gut microbiota.

As an approximation of a stability landscape, the energy landscape would provide a reliable map for those who seek an effective path between different community compositions. In conservation biology, knowledge of the paths by which communities are assembled helps ecologists to understand the role of history in shaping current communities, and is important for effective community restoration (Weiher and Keddy 1999, Lockwood & Samuels 2004, Suding et al. 2004, Wilsey et al. 2015, Young et al. 2005, 2015). In other words, when historical contingency occurs, restoring and maintaining native biodiversity may require specific sequences of exotic species removal and/or native species introduction. This is also relevant to agriculture, e.g., the successful inoculation of agricultural soils with beneficial fungi or other microbes may depend on the timing of inoculation relative to plant growth, as well as the profile of other soil microbes (Verbruggen et al. 2013, Toju et al. 2018). In medicine, the relevance of historical contingency in community assemblies to curing some human diseases is being recognized (Costello et al. 2012, Fierer et al. 2012, Lam & Monack 2014, Devevey et al. 2015). Clinically meaningful evidence for the potential application of modulating the intestinal microbiota for therapeutic gain has created considerable interest and enthusiasm (Smits et al. 2013, Li et al. 2016, Shetty et al. 2017). In addition to *Clostridium difficile* Infection (CDI), a disruption to the gut microbiota is associated with, e.g., irritable bowel syndrome, autism, obesity, and cavernous cerebral malformations (Karczewski et al. 2014, Cox et al. 2015, Tang et al. 2017). Driving disrupted microbial communities back to their healthy states could offer novel solutions to prevent and treat complex human diseases.

There are alternative methodologies to study stability landscapes. Dynamical models (such as differential equations) are able to describe shifts in community compositions based on the gradual changes in population abundance of species (Gravel et al. 2011). If we are able to estimate the parameters of a dynamical model, it allows us to directly study the stability landscape that is inferred from observational data. However, it is generally difficult to develop fully mechanistic models for multi-species communities. The complex nature of microbial interactions makes it difficult to formulate all the present relationships into mathematical formulations (Hartig and Dormann 2013; Perretti et al. 2013a, b; De Angelis and Yurek 2015). Moreover, a theoretical study proved that finding a precise dynamical equation for a time-series is, in general, computationally intractable even with any amount/quality of data (Cubitt et al. 2012). Another potential approach may be developing a Markov chain model with a transition matrix between different community compositions (Wootton 2001). This can be done if we can obtain multiple observations on two consecutive community compositions. However, considering the number of possible paths in a multispecies system, it might not be a realistic approach. Actually, in our LV model with a constant environment, among randomly sampled 128 pairs of consecutive community compositions (comparative to the 256 data points used to fit pairwise maximum entropy models), 88.1% (s.d. 3.8) of them were observed only once. Thus, it is obvious that it requires much more data to reconstruct a transition matrix. Recently developed methods for reconstructing low dimensional potential landscape (Gibson et al. 2017, Shaw et al. 2019, Chang et al. 2019) would be an appealing option, but since this approach projects microbial dynamics onto a low dimensional potential landscape, it is unable to reproduce overall compositional stability as we see in this paper. Also, it is unable to track the change of compositional stability across environmental parameters. Considering the data set size and observational procedure required, our method is a realistic option to study the compositional stability of empirical communities.

There are still some flaws that need to be addressed. First, greater sophistication of the model fitting framework, including sparse modelling, would improve the performance of our approach. Second, further verification is required to assess the effect of replacing species’ abundance with presence/absence status. Does the pairwise maximum model always provide good approximation to the stability landscape? For example, since our approach relies on a gradient system, it cannot account for attractor dynamics that often appear in ecological systems. Heteroclinic cycles that cause cyclic alteration of community compositions (Morton and Law 1997, Fukami et al. 2015) are one example. These dynamics may still be identified as a set of stable states separated by low energy barriers, though the overall consequence of approximating dynamics in a continuous phase space into a coarse-grained phase space (where nodes of the weighted network represent each sub-system of the original phase space) is unknown. Finally, causal relationships between species’ presence/absence status and the transition of one community state to another are not well represented using our approach. Incorporating causal analysis (e.g., Sugihara et al. 2012, Runge et al. 2017) will strengthen our approach, especially when considering applications to control community states.

## Conclusion

We have demonstrated the effectiveness of pairwise maximum entropy models as an approximation of overall compositional stability, which we defined as a stability landscape, of multispecies communities in a changing environment. The framework of energy landscape analysis played a prominent role in the success of this approach. Our model opens up new research directions encompassing the concept of alternative stability in community assemblies of multispecies communities, and the change in dynamical stability across environmental gradients, which have mainly been studied in low-dimensional systems in the past decades. There is urgent need for a methodology that is able to account for compositional dynamics in multispecies communities. Although some further verification and improvement is required, we believe that the methodological advancement presented here will be a new systemic paradigm for developing a predictive theory for real-world ecological communities (Mouquet et al. 2015).

## Acknowledgements

We would like to thank two anonymous reviewers, and Sammuel R.P.-J. Ross, Taku Kadoya and Hirokazu Toju. This work was supported by the Management Expenses Grant for RIKEN BioResource Research Center, MEXT, and in part by the Center of Innovation Program from Japan Science and Technology Agency (JST) (to K.S. and S.N.), JST PRESTO JPMJPR16E9 (to S.N.), JPMJPR1537 (to S.F.), JST ERATO JPMJER1902 (to S.F.), the Japan Society for the Promotion of Science (JSPS) KAKENHI JP20K06820 and JP20H03010 (to K.S.), JP15H05707 (to S.N.), JP18H04805 (to S.F.), AMED-CREST JP20gm1010009 (to S.F.), the Takeda Science Foundation (to S.F.), the Food Science Institute Foundation (to S.F.) and the Program for the Advancement of Research in Core Projects under Keio University’s Longevity Initiative (to S.F.). We declare that we have no conflict of interest.

## Appendix S1 Parameter fitting

The maximum likelihood estimate for the model parameters can be obtained by minimizing the discrepancy between the values of the data’s sufficient statistics and the corresponding sufficient statistics within the model (Bickel and Doksum 1977, Azaele et al. 2010, Murphy 2012, Harris 2015, Lee and Hastie 2015).

### Gradient descent algorithm

For a dataset without observation on environmental conditions, i.e., whose energy is given by eq. (4), a gradient descent algorithm can be applied (Watanabe et al. 2014a,b, Harris 2015, Harris 2016). For a model with parameter *h*\* and *J*\*, let the expected probability of species i be 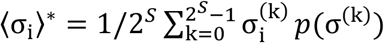 and the co-occurrence be 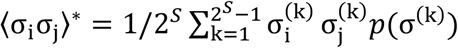. The parameter *h* and *J* can be fitted to the data by iteratively adjusting 〈σ_i_〉* and 〈σ_i_σ_j_〉* toward the mean occurrence and co-occurrence calculated from the observational data, 〈σ_i_〉 and 〈σ_i_σ_j_〉. Here, the parameters are updated as at

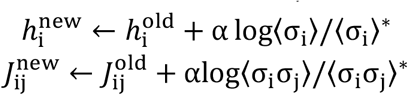

each step. We set the learning rate α = 0.25 and the maximum number of iterations *T* = 5000, according to the preliminary analysis where we checked the convergence of model parameters.

### Stochastic approximation

The likelihood function of the pairwise maximum entropy model becomes computationally intractable when we need to include environmental condition (as in eq. (2)), because it requires repeating the above computations independently for every sample. Therefore, it calls for a different model-fitting algorithm. Here, following Harris (2015), we introduce a *stochastic approximation* (Robbins and Monro 1951, Salakhutdinov and Hinton 2012) for this purpose. This algorithm replaces the intractable computations with tractable Monte Carlo estimates of the same quantities. Despite the sampling error introduced by this substitution, stochastic approximation provides strong guarantees for eventual convergence to the maximum likelihood estimate (Younes 1999, Salakhutdinov and Hinton 2012).

Stochastic approximation (Robbins and Monro 1951, Salakhutdinov and Hinton 2012) estimates the expected values of the sufficient statistics by averaging over a more manageable number of simulated assemblages during each model-fitting iteration, while still retaining maximum likelihood convergence. The advantage of this algorithm is that *Z* (eq. (3)) does not have to be calculated at each step, which significantly improves computational efficiency. This is due to the use of a heat-bath algorithm that only requires calculating energy of two adjacent community compositions. The procedure iterates the following steps:

1. Set *t* = 0, initial learning rate α_0_ = 0.1, logistic priors as *p_h_* = −tanh (*h*/2/2)/2, *p_g_* = −tanh (*g*/2/2)/2 and *p_J_* = −tanh (*J*/0.5/2)/2 and initialize parameter values for *h,J, g*, and the expected sample states X*(0) = X. We set Y* = Y throughout the calculation.
2. Calculate learning rate α as:

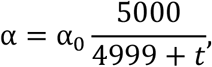

momentum *m* as:

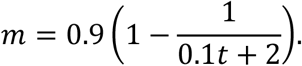
3. For 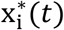 from i = 1 to *N*, run one step heat-bath algorithm based on current parameters (*h, J* and *g*): transition from the current community composition σ^(k)^ to one of its *S* adjacent community composition σ^(k′)^, selected with probability 1/*S*, was attempted (σ^(k)^ and σ^(k′)^ differs only with respect to the presence/absence status of one of *S* species). The transition to the selected state took place with probability e^−*E*(σ(k′)|ε(i))^/(e^−*E*(σ(k)|ε(i))^ + e^−*E*(σ(k′)|ε(i))^). Here e^−*E*(σ(k)|ε(i))^ and e^−*E*(σ(k′)|ε(i))^ occurs, the sample state is updated as 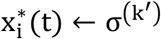.
4. Subtract the simulated sufficient statistics from the observed ones to calculate the approximate likelihood gradient. Sufficient statistics are calculated as, 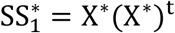 (here, (X*)^t^ is the transpose of X*), and 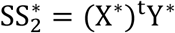. Then, we obtain the difference of sufficient statistics as:

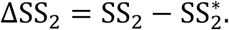

and

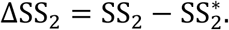 Here, SS_1_ and SS_2_ is the corresponding sufficient statistics calculated from actual data (i.e., SS_1_ = XX^t^ and SS_2_ = X^t^ Y).
5. Adjust the model parameters to climb the approximate gradient, using a schedule of step sizes as:

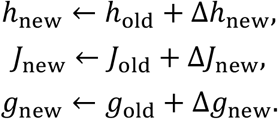 Here,

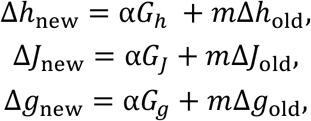

and,

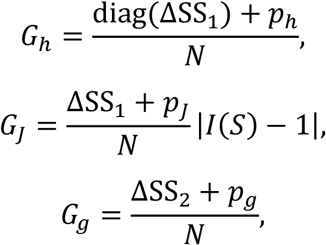

are the approximated likelihood gradients. Here, *I(S)* is a *S* × *S* identity matrix.
6. Set *h*_new_, *J*_new_, *g*_new_, Δ*h*_new_; Δ*J*_new_ and Δ*g*_new_ as *h*_old_, *J*_old_, *g*_old_, Δ*h*_old_, Δ*J*_old_ and Δ*g*_old_, respectively. If *t* < *T*, increment t by 1 and back to 2, else terminate the loop.

The simulations in Step 3 use a one step heat-bath algorithm (Gibbs sampling) to generate a community composition distribution based on the model’s current parameter estimates. While the subsequent community compositions produced by Gibbs sampling are autocorrelated, this does not prevent convergence to the maximum likelihood estimates (Younes 1999, Salakhutdinov and Hinton 2012). Approximated likelihood gradients in Step 5 match those of gradient descent, except that they are averaged over a set of Monte Carlo samples rather than over all possible community composition. These gradients were augmented with a momentum term (Hinton 2012) and by regularizers based on a logistic prior with location 0 and scale 2.0 (for environmental responses) or 0.5 (for pairwise relationships). We set hyperparameters in this algorithm, including a maximum number of iteration steps *T* = 50000, according to the preliminary analysis where we checked the convergence of model parameters.

## Appendix S2 Parameter values and models used to generate data sets

The interaction matrix used to generate a data set in *Analysis of a competitive Lotka-Volterra system, Energy landscape of the LV system, Emulating community assembly dynamics*, are in Table S1.

The response vector *b* in *Energy landscape across environmental gradient* is:

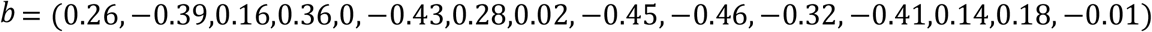

In *Benchmarking*, the interaction matrix *A* was generated so that the connectance *c_A_* = 0.5, and its non-zero diagonal elements (interspecific competition) were drawn from a normal distribution with a mean *μ_A_* = 0.6 and variance σ_*A*_ = 0.2. We fixed the diagonal elements (intraspecific interaction) as *a_ii_* = 1. To obtain the result under type II functional response (G in Fig. 7), we extended the eq.(5) as

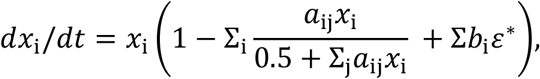

and to obtain data set with noise (H in Fig. 7), we used,

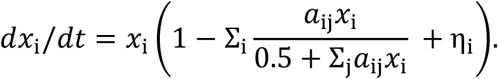

Here, η_i_ is assumed to be the i.i.d noise drawn from a normal distribution of mean 0 and s.d. 0.1.

**Table S1.** Elements of matrix *A*.

|  |  |  |  |  |  |  |  |  |  |  |  |  |  |  |  |
| --- | --- | --- | --- | --- | --- | --- | --- | --- | --- | --- | --- | --- | --- | --- | --- |
| 1 | 0 | 0.5<br>8 | 0.6<br>2 | 0 | 0.6<br>1 | 0.2<br>9 | 0 | 0 | 0.5<br>5 | 0.6<br>9 | 0.5<br>4 | 0.6<br>6 | 0 | 0 | 0 |
| 0.8 | 1 | 0 | 0.5<br>3 | 0.6<br>1 | 0.6<br>5 | 0 | 0.7<br>8 | 0 | 0.5<br>8 | 0.7<br>4 | 0.7<br>2 | 0 | 0.5<br>2 | 0.6<br>2 | 0 |
| 0 | 0 | 1 | 0.6<br>1 | 0.6<br>8 | 0 | 0 | 0 | 0 | 0 | 0 | 0.4<br>5 | 0.6<br>2 | 0.6<br>7 | 0.7 | 0 |
| 0.3<br>3 | 0.5<br>6 | 0.5<br>8 | 1 | 0.5<br>1 | 0.5 | 0.6<br>5 | 0.5<br>2 | 0 | 0.5<br>5 | 0 | 0 | 0.6<br>1 | 0 | 0.5<br>4 | 0.4<br>8 |
| 0.7<br>7 | 0 | 0.6<br>3 | 0.5<br>7 | 1 | 0.6<br>7 | 0.5<br>5 | 0.6<br>1 | 0 | 0.6<br>3 | 0 | 0.5<br>7 | 0 | 0.4<br>8 | 0.5<br>2 | 0.8<br>6 |
| 0 | 0 | 0.5<br>8 | 0.4<br>3 | 0 | 1 | 0.4<br>7 | 0.7<br>5 | 0.5<br>3 | 0 | 0.4<br>9 | 0.4<br>9 | 0.4<br>5 | 0 | 0.5<br>1 | 0 |
| 0.5<br>4 | 0 | 0 | 0.6<br>1 | 0.7 | 0 | 1 | 0 | 0.4 | 0 | 0.6<br>3 | 0 | 0.7<br>9 | 0.5<br>5 | 0 | 0 |
| 0 | 0 | 0.6 | 0.5<br>6 | 0.4<br>3 | 0 | 0.7<br>6 | 1 | 0 | 0 | 0.5<br>1 | 0.9 | 0.5<br>9 | 0.7<br>7 | 0.5<br>2 | 0 |
| 0.5<br>5 | 0 | 0 | 0.6<br>6 | 0.6<br>3 | 0 | 0.6<br>1 | 0 | 1 | 0 | 0.4<br>4 | 0 | 0 | 0 | 0.4<br>8 | 0.3<br>9 |
| 0 | 0.3<br>8 | 0.5<br>6 | 0.5<br>5 | 0 | 0.4<br>7 | 0.5<br>6 | 0.7<br>6 | 0 | 1 | 0 | 0 | 0 | 0.5<br>4 | 0.6<br>2 | 0 |
| 0.4<br>4 | 0 | 0.4<br>9 | 0 | 0.7<br>1 | 0.5<br>2 | 0 | 0.4<br>7 | 0.7<br>7 | 0 | 1 | 0.7<br>5 | 0.8<br>4 | 0 | 0.6<br>7 | 0.7<br>5 |
| 0.5<br>8 | 0.3<br>3 | 0 | 0.4<br>7 | 0.6<br>8 | 0.4<br>6 | 0 | 0.4<br>2 | 0.6<br>1 | 0.6 | 0.6<br>1 | 1 | 0.6<br>3 | 0.7 | 0.6<br>9 | 0 |
| 0.7 | 0 | 0 | 0.5<br>7 | 0 | 0 | 0.6<br>2 | 0 | 0 | 0.7<br>8 | 0.6<br>3 | 0.6<br>7 | 1 | 0 | 0 | 0.5<br>1 |
| 0.5<br>4 | 0.7<br>4 | 0 | 0 | 0.5<br>1 | 0 | 0.7<br>8 | 0.4<br>9 | 0 | 0 | 0.7<br>1 | 0.4<br>4 | 0.4<br>3 | 1 | 0.6<br>4 | 0 |
| 0.4<br>5 | 0 | 0.6<br>2 | 0 | 0.5<br>4 | 0.7<br>3 | 0 | 0.4<br>7 | 0.6<br>7 | 0.7<br>7 | 0 | 0.7 | 0.6<br>9 | 0.5<br>3 | 1 | 0 |
| 0.8<br>1 | 0.4<br>6 | 0 | 0.6<br>7 | 0.3<br>8 | 0.6<br>5 | 0 | 0 | 0.5<br>5 | 0.8<br>5 | 0.7<br>6 | 0 | 0.5<br>8 | 0 | 0.5<br>5 | 1 |

## Glossary of terms

Actual features: features characterizing the stability landscape and is calculated by LV data set; these features are comparable to the features of an energy landscape.
Basin of attraction: in an energy landscape, defined as a set of community compositions that reach one distinct stable state when assembly processes are completely deterministic; in LV data set, it is identified by a stable state to which a community composition most frequently converged (if there is more than one such state, it belongs to all of them).
Effective boundary: community compositions in emulated compositional dynamics having the highest energy during the transition from one stable state to another.
Empirical probability: one of actual features; the ratio of the number of observations of σ^(k)^ to the total number of observations.
Emulated compositional dynamics: compositional dynamics constrained by an energy landscape; it is generated by using the heat-bath (also known as Gibbs sampling) method.
Energy barrier: the energy level that need to go up during the transition from one stable state to another.
Energy landscape: a weighted network whose nodes represent unique community compositions and links represent transition path between them; nodes are weighted according to energy E given by eq.(2) or (4); an energy landscape is the approximation of a stability landscape based on the maximum entropy principle given observational data; energy landscape analysis is the analysis of topological and connection attributes of an energy landscape.
Energy minima: community compositions having the lowest energy compared to all neighboring compositions, and thus constitute end-points when assembly processes are completely deterministic (i.e., when transition of community compositions always go down the energy landscape); we identify energy minima of an energy landscape as stable states of a stability landscape.
Extended pairwise maximum entropy model: an extension of the pairwise maximum entropy model (Markov network) including a term representing environmental effects. We referred the two models as the pairwise maximum entropy models all together.
Imbalance score (IS): one of actual features; quantifies how stable states to which a community composition in LV data set converges are uniquely determined.
LV data set: a data set generated by the LV competition model; it is used to calculate actual features.
Relative convergence time (RCT): one of actual features; the average number of different community compositions that a community composition undergoes before converging to a stable state; it is normalized to have a value between 0 and 1 and indicates distance from a community composition to a stable state to which it converges.
Rescaled energy: the energy of a community composition normalized to take 0 at a stable state and 1 at the community composition that has the highest energy within an attractive basin; for σ^(k)^ it is calculated as, 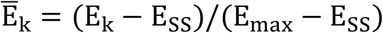 where E_k_ is the energy of σ^(k)^, E_SS_ is the energy of stable state to which basin σ^(k)^ belongs, and E_max_ is the energy of a community composition that is the highest within the basin of attraction.
Stable state: given a fixed set of species, a community composition that can be an end state of assembly sequences.
Stability landscape: a structure that governs the overall compositional stability of an ecological community; it can be represented as a graph with a set of community compositions and transition paths between them (Figure 1).
Tipping point: the community composition located at the lowest part of the ridge between two basins of attraction.
Transition channels: a fraction of effective boundary that mediates most of the transition between stable states.

## References

Araújo, M. B., Luoto M. (2007). The importance of biotic interactions for modelling species distributions under climate change. Global Ecology and Biogeography, 16, 743–753.

Azaele, S., Muneepeerakul, R., Rinaldo, A., Rodriguez-Iturbe, I. (2010). Inferring plant ecosystem organization from species occurrences. Journal of theoretical biology, 262(2), 323–329.

Bakken, J. S., Borody, T., Brandt, L. J., Brill, J. V., Demarco, D. C., Franzos, M. A., et al. (2011). Treating Clostridium difficile infection with fecal microbiota transplantation. Clinical Gastroenterology and Hepatology, 9(12), 1044–1049.

Barbaro, L., Allan, E., Ampoorter, E., Castagneyrol, B., Charbonnier, Y., De Wandeler, H., et al. (2019). Biotic predictors complement models of bat and bird responses to climate and tree diversity in European forests. Proceedings of the Royal Society B, 286(1894), 20182193.

Barner, A. K., Coblentz, K. E., Hacker, S. D., Menge, B. A. (2018). Fundamental contradictions among observational and experimental estimates of non-trophic species interactions. Ecology, 99(3), 557–566.

Becker, O. M., Karplus, M. (1997). The topology of multidimensional potential energy surfaces: Theory and application to peptide structure and kinetics. The Journal of chemical physics, 106(4), 1495–1517.

Beisner, B. E., Haydon, D. T. Cuddington, K. Alternative stable states in ecology. Frontiers in Ecology and the Environment, 1, 376–382 (2003).

Bolyen, E., Rideout, J. R., Dillon, M. R., Bokulich, N. A., Abnet, C. C., Al-Ghalith, G. A., et al. (2019). Reproducible, interactive, scalable and extensible microbiome data science using QIIME 2. Nature biotechnology, 37(8), 852–857.

Britton, R. A., Young, V. B. (2014). Role of the intestinal microbiota in resistance to colonization by Clostridium difficile. Gastroenterology, 146(6), 1547–1553.

Capitán, J. A., Cuesta, J. A., Bascompte, J. (2011). Statistical mechanics of ecosystem assembly. Physical Review Letters, 103(16), 168101.

Caporaso, J. G., Kuczynski, J., Stombaugh, J., Bittinger, K., Bushman, F. D., Costello, E. K., et al. (2010). QIIME allows analysis of high-throughput community sequencing data. Nature methods, 7(5), 335–336.

Carding, S., Verbeke, K., Vipond, D. T., Corfe, B. M., Owen, L. J. (2015). Dysbiosis of the gut microbiota in disease. Microbial ecology in health and disease, 26(1), 26191.

Castledine, M., Sierocinski, P., Padfield, D., Buckling, A. (2020). Community coalescence: an eco-evolutionary perspective. Philosophical Transactions of the Royal Society B, 375(1798), 20190252.

Chang, W. K., VanInsberghe, D., Kelly, L. (2020). Towards a potential landscape framework of microbiome dynamics. bioRxiv, 584201.

Chase, J.M. (2014). Spatial scale resolves the niche versus neutral theory debate. Journal of vegetation science, 25, 319–322.

Clark, N. J., Wells, K., Lindberg, O. (2018). Unravelling changing interspecific interactions across environmental gradients using Markov random fields. Ecology, 99(6), 1277–1283.

Costello, E. K., Stagaman, K., Dethlefsen, L., Bohannan, B. J., Relman, D. A. (2012). The application of ecological theory toward an understanding of the human microbiome. Science, 336(6086), 1255–1262.

Cox, L. M., Blaser, M. J. (2015). Antibiotics in early life and obesity. Nature Reviews Endocrinology, 11(3), 182.

Cubitt, T. S., Eisert, J., Wolf, M. M. (2012). Extracting dynamical equations from experimental data is NP hard. Physical Review Letters, 108, 120503.

D’Amen, M., Mod, H. K., Gotelli, N. J., Guisan, A. (2018). Disentangling biotic interactions, environmental filters, and dispersal limitation as drivers of species co-occurrence. Ecography, 41, 1233–1244.

DeAngelis, D. L., Yurek, S. (2015). Equation-free modeling unravels the behavior of complex ecological systems. Proceedings of the National Academy of Sciences, 112, 3856–3857.

Devevey, G., Dang, T., Graves, C. J., Murray, S., Brisson, D. (2015). First arrived takes all: inhibitory priority effects dominate competition between co-infecting Borrelia burgdorferi strains. BMC microbiology, 15(1), 61.

Ding, T., Schloss, P. D. (2014). Dynamics and associations of microbial community types across the human body. Nature, 509(7500), 357–360.

Drake, J. A. (1991). Community-assembly mechanics and the structure of an experimental species ensemble. The American Naturalist, 137(1), 1–26.

Elith, J., H. Graham, C., P. Anderson, R., Dudík, M., Ferrier, S., Guisan, A., et al. (2006). Novel methods improve prediction of species’ distributions from occurrence data. Ecography, 29(2), 129–151.

Ezaki, T., Sakaki, M., Watanabe, T., Masuda, N. (2018). Age-related changes in the ease of dynamical transitions in human brain activity. Human brain mapping, 39(6), 2673–2688.

Ezaki, T., Watanabe, T., Ohzeki, M., Masuda, N. (2017). Energy landscape analysis of neuroimaging data. Philosophical Transactions of the Royal Society A: Mathematical, Physical and Engineering Sciences, 375(2096), 20160287.

Fierer, N., Ferrenberg, S., Flores, G. E., González, A., Kueneman, J., Legg, T., et al. (2012). From animalcules to an ecosystem: application of ecological concepts to the human microbiome. Annual Review of Ecology, Evolution, and Systematics, 43, 137–155.

Franklin, J. (2010). Mapping species distributions: spatial inference and prediction. Cambridge University Press.

Freilich, M. A., Wieters, E., Broitman, B. R., Marquet, P. A., Navarrete, S. A. (2018). Species co-occurrence networks: Can they reveal trophic and non-trophic interactions in ecological communities? Ecology, 99(3), 690–699.

Fukami, T. (2010). Community assembly dynamics in space. In Community ecology: Processes, models, and applications. Edited by Herman A. Verhoef and Peter J. Morin, 45–54. Oxford: Oxford Univ. Press.

Fukami, T. (2015). Historical contingency in community assembly: integrating niches, species pools, and priority effects. Annual Review of Ecology, Evolution, and Systematics, 46, 1–23.

Fukami, T., Dickie, I. A., Paula Wilkie, J., Paulus, B. C., Park, D., Roberts, A., et al. (2010). Assembly history dictates ecosystem functioning: evidence from wood decomposer communities. Ecology letters, 13(6), 675–684.

Fukami, T., Morin, P. J. (2003). Productivity-biodiversity relationships depend on the history of community assembly. Nature, 424(6947), 423.

Gerber, G. K. (2014). The dynamic microbiome. FEBS letters, 588(22), 4131–4139.

Gibson, T. E., Carey, V., Bashan, A., Hohmann, E. L., Weiss, S. T., Liu, Y. Y. (2017). On the Stability Landscape of the Human Gut Microbiome: Implications for Microbiome-based Therapies. bioRxiv, 176941.

Gilks, W. R., Richardson, S., Spiegelhalter, D. J. (1996). Introducing Markov chain Monte Carlo. Markov chain Monte Carlo in practice. Chapman & Hall.

Gilpin, M. E., Case, T. J. (1976). Multiple domains of attraction in competition communities. Nature, 261(5555), 40–42.

Gravel, D., Guichard, F., Hochberg, M. E. (2011). Species coexistence in a variable world. Ecology letters, 14(8), 828–839.

Harris, D. J. (2015). Multi-Process Statistical Modeling of Species’ Joint Distributions. University of California, Davis.

Harris, D. J. (2016). Inferring species interactions from co-occurrence data with Markov networks. Ecology, 97(12), 3308–3314.

Harte, J. (2011). Maximum entropy and ecology: a theory of abundance, distribution, and energetics. OUP Oxford.

Harte, J., Newman, E. A. (2014). Maximum information entropy: a foundation for ecological theory. Trends in ecology & evolution, 29(7), 384–389.

Hartig, F. Dormann, C. F. (2013). Does model-free forecasting really outperform the true model? Proceedings of the National Academy of Science, 110, E3975–E3975.

Holling, C. S. (1996). Engineering resilience versus ecological resilience. In Engineering within Ecological Constraints, ed. PC Schulze, pp. 31–44. Washington, DC: Natl. Acad. Press.

Jaynes, E.T., (1982). On the rationale of maximum-entropy methods. Proceedings of the IEEE, 70, 939–952.

Jiang, L., Joshi, H., Flakes, S. K., Jung, Y. (2011). Alternative community compositional and dynamical states: the dual consequences of assembly history. Journal of Animal Ecology, 80(3), 577–585.

Kadowaki, K., Inouye, B. D., Miller T. E. (2012). Assembly-History Dynamics of a Pitcher-Plant Protozoan Community in Experimental Microcosms. PLoS One, 7(8), e42651.

Karczewski, J., Poniedziałek, B., Adamski, Z., Rzymski, P. (2014). The effects of the microbiota on the host immune system. Autoimmunity, 47(8), 494–504.

Kelly C. P., LaMont J. T. (2008). *Clostridium difficile—*more difficult than ever. New England Journal of Medicine, 359:1932–1940, 2008.

Keymer, J. E., Marquet, P. A., Velasco-Hernández, J. X., Levin, S. A. (2000). Extinction thresholds and metapopulation persistence in dynamic landscapes. The American Naturalist, 156(5), 478–494.

Kissling, W. D., Dormann, C. F., Groeneveld, J., Hickler, T., Kühn, I., McInerny, G. J., et al. (2012). Towards novel approaches to modelling biotic interactions in multispecies assemblages at large spatial extents. Journal of Biogeography, 39(12), 2163–2178.

Lahti, L., Salojärvi, J., Salonen, A., Scheffer, M., De Vos, W. M. (2014). Tipping elements in the human intestinal ecosystem. Nature communications, 5(1), 1–10.

Lam, L. H., Monack, D. M. (2014). Intraspecies competition for niches in the distal gut dictate transmission during persistent Salmonella infection. PLoS Pathogens, 10(12), e1004527.

Langille, M. G., Meehan, C. J., Koenig, J. E., Dhanani, A. S., Rose, R. A., Howlett, S. E., et al. (2014). Microbial shifts in the aging mouse gut. Microbiome, 2(1), 50.

Law, R., A. J. Weatherby, P. H. Warren. (2000). On the invasibility of persistent protist communities. Oikos, 88, 319–326.

Law, R., Morton, R. D. (1993). Alternative permanent states of ecological communities. Ecology, 74(5), 1347–1361.

Leach, K., Montgomery, W. I., Reid, N. (2016). Modelling the influence of biotic factors on species distribution patterns. Ecological Modelling, 337, 96–106.

Legendre, P., Mi, X., Ren, H., Ma, K., Yu, M., Sun, I.F., et al. (2009). Partitioning beta diversity in a subtropical broad-leaved forest of China. Ecology, 90, 663–674.

Li, S. S., Zhu, A., Benes, V., Costea, P. I., Hercog, R., Hildebrand, F., et al. (2016). Durable coexistence of donor and recipient strains after fecal microbiota transplantation. Science 352, 586–586.

Lockwood, J. L., Samuels, C. L. (2004). Assembly models and the practice of restoration. In Assembly Rules and Restoration Ecology: Bridging the Gap Between Theory and Practice, ed. VM Temperton, RJ Hobbs, T Nuttle, S Halle, pp. 55–70. Washington, DC: Island Press.

May, R. M. (1977). Thresholds and breakpoints in ecosystems with a multiplicity of stable states. Nature, 269(5628), 471–477.

Meier, E. S., Kienast, F., Pearman, P. B., Svenning, J. C., Thuiller, W., Araújo, M. B., et al. (2010). Biotic and abiotic variables show little redundancy in explaining tree species distributions. Ecography, 33(6), 1038–1048.

Mertes, K., Jetz, W. (2018). Disentangling scale dependencies in species environmental niches and distributions. Ecography, 41, 1604–1615.

Meynard, C.N., Lavergne, S., Boulangeat, I., Garraud, L., Van Es, J., Mouquet, N., et al. (2013). Disentangling the drivers of metacommunity structure across spatial scales. Journal of biogeography, 40, 1560–1571.

Morton, R. D., Law, R. (1997). Regional species pools and the assembly of local ecological communities. Journal of Theoretical Biology, 187, 321–331.

Mouquet, N., Lagadeuc, Y., Devictor, V., Doyen, L., Duputié, A., Eveillard, D., et al. (2015). Predictive ecology in a changing world. Journal of Applied Ecology, 52(5), 1293–1310.

Mueller, U. G., Sachs, J. L. (2015). Engineering microbiomes to improve plant and animal health. Trends in microbiology, 23(10), 606–617.

Nakanishi, Y., Nozu, R., Ueno, M., Hioki, K., Ishii, C., Murakami, S., et al. (2020). Longitudinal analyses reveal that aging-related alterations in the intestinal environment lead to gut dysbiosis with the potential to induce obesity. Research Square [https://doi.org/10.21203/rs.3.rs-119480/v1].

Nguyen, H. C., Zecchina, R., Berg, J. (2017). Inverse statistical problems: from the inverse Ising problem to data science. Advances in Physics, 66(3), 197–261.

Ockendon, N., Baker, D. J., Carr, J. A., White, E. C., Almond, R. E., Amano, T., et al. (2014). Mechanisms underpinning climatic impacts on natural populations: altered species interactions are more important than direct effects. Global change biology, 20(7), 2221–2229.

Parisien, M. A., Moritz, M. A. (2009). Environmental controls on the distribution of wildfire at multiple spatial scales. Ecological Monographs, 79(1), 127–154.

Perretti, C. T., Munch, S. B., Sugihara, G. (2013a) Model-free forecasting outperforms the correct mechanistic model for simulated and experimental data. Proceedings of the National Academy of Science, 110, 5253–5257.

Perretti, C. T., Munch, S. B., Sugihara, G. (2013b). Reply to Hartig and Dormann: the true model myth. Proceedings of the National Academy of Science, 110, E3976–E3977.

Phillips, S. J., Anderson, R. P., Schapire, R. E. (2006). Maximum entropy modeling of species geographic distributions. Ecological modelling, 190(3-4), 231–259.

Phillips, S. J., Dudík, M., Schapire, R. E. (2004). A maximum entropy approach to species distribution modeling. In Proceedings of the twenty-first international conference on Machine learning (p. 83).

Pu, Z., Jiang, L. (2015). Dispersal among local communities does not reduce historical contingencies during metacommunity assembly. Oikos, 124(10), 1327–1336.

Rillig, M. C., Antonovics, J., Caruso, T., Lehmann, A., Powell, J. R., Veresoglou, S. D., et al. (2015). Interchange of entire communities: microbial community coalescence. Trends in Ecology and Evolution, 30(8), 470–476.

Runge, J., Nowack, P., Kretschmer, M., Flaxman, S., Sejdinovic, D. (2019). Detecting and quantifying causal associations in large nonlinear time series datasets. Science Advances, 5(11), eaau4996.

Sbahi, H., Di Palma, J. A. (2016). Faecal microbiota transplantation: applications and limitations in treating gastrointestinal disorders. BMJ open gastroenterology, 3(1).

Scheffer, M., Carpenter, S. R., Dakos, V., van Nes, E. H. (2015). Generic indicators of ecological resilience: inferring the chance of a critical transition. Annual Review of Ecology, Evolution, and Systematics, 46, 145–167.

Scheffer, M., Carpenter, S. R., Foley, J. A., Folke, C., Walker, B. (2001). Catastrophic shifts in ecosystems. Nature, 413(6856), 591.

Scheffer, M., Carpenter, S. R., Lenton, T. M., Bascompte, J., Brock, W., Dakos, V., et al. (2012). Anticipating critical transitions. Science, 338(6105), 344–348.

Scheffer, M., Jeppesen, E. (2007). Regime shifts in shallow lakes. Ecosystems, 10(1), 1–3.

Schneidman, E., Berry, M. J., Segev, R., Bialek, W. (2006). Weak pairwise correlations imply strongly correlated network states in a neural population. Nature, 440(7087), 1007–1012.

Schröder, A., Persson, L., De Roos, A. M. (2005). Direct experimental evidence for alternative stable states: a review. Oikos, 110(1), 3–19.

Shaw, L. P., Bassam, H., Barnes, C. P., Walker, A. S., Klein, N., Balloux, F. (2019). Modelling microbiome recovery after antibiotics using a stability landscape framework. The ISME journal, 13(7), 1845–1856.

Shetty, S. A., Hugenholtz, F., Lahti, L., Smidt, H., de Vos, W. M. (2017). Intestinal microbiome landscaping: insight in community assemblage and implications for microbial modulation strategies. FEMS microbiology reviews, 41(2), 182–199.

Shipley, B., Vile, D., & Garnier, É. (2006). From plant traits to plant communities: a statistical mechanistic approach to biodiversity. Science, 314(5800), 812–814.

Smits, L. P., Bouter, K. E., de Vos, W. M. et al. (2013). Therapeutic potential of fecal microbiota transplantation. Gastroenterology 145:946–53.

Sommer, F., Anderson, J. M., Bharti, R., Raes, J., Rosenstiel, P. (2017). The resilience of the intestinal microbiota influences health and disease. Nature Reviews Microbiology, 15(10), 630–638.

Staniczenko, P. P., Sivasubramaniam, P., Suttle, K. B., Pearson, R. G. (2017). Linking macroecology and community ecology: refining predictions of species distributions using biotic interaction networks. Ecology letters, 20(6), 693–707.

Suding, K. N., Gross, K. L., Houseman, G. R. (2004). Alternative states and positive feedbacks in restoration ecology. Trends in Ecology and Evolution, 19, 46–53.

Sugihara, G., May, R., Ye, H., Hsieh, C. H., Deyle, E., Fogarty, M., et al. (2012). Detecting causality in complex ecosystems. Science, 338(6106), 496–500.

Tang, A. T., Choi, J. P., Kotzin, J. J., Yang, Y., Hong, C. C., Hobson, N., et al. (2017). Endothelial TLR4 and the microbiome drive cerebral cavernous malformations. Nature, 545(7654), 305.

Thompson, L. R., Sanders, J. G., McDonald, D., Amir, A., Ladau, J., Locey, K. J., et al. (2017). A communal catalogue reveals Earth’s multiscale microbial diversity. Nature, 551(7681), 457–463.

Thuiller, W., Pollock, L. J., Gueguen, M., Münkemüller, T. (2015). From species distributions to meta-communities. Ecology letters, 18(12), 1321–1328.

Toju, H., Abe, M. S., Ishii, C., Hori, Y., Fujita, H., Fukuda, S. (2020). Scoring Species for Synthetic Community Design: Network Analyses of Functional Core Microbiomes. Frontiers in microbiology, 11, 1361.

Toju, H., Peay, K. G., Yamamichi, M., Narisawa, K., Hiruma, K., Naito, K., et al. (2018). Core microbiomes for sustainable agroecosystems. Nature Plants, 4(5), 247–257.

Verbruggen, E., van der Heijden, M. G., Rillig, M. C., Kiers, E. T. (2013). Mycorrhizal fungal establishment in agricultural soils: factors determining inoculation success. New Phytologist, 197, 1104–1109.

Viana, D.S., Chase, J.M. (2019). Spatial scale modulates the inference of metacommunity assembly processes. Ecology, 100, e02576.

Wales, D. J., Miller, M. A., Walsh, T. R. (1998). Archetypal energy landscapes. Nature, 394(6695), 758.

Walker, B., Holling, C. S., Carpenter, S., Kinzig, A. (2004). Resilience, adaptability and transformability in social–ecological systems. Ecology and society, 9(2).

Warren, P. H., Law, R., Weatherby, A. J. (2003). Mapping the assembly of protist communities in microcosms. Ecology, 84(4), 1001–1011.

Watanabe, T., Hirose, S., Wada, H., Imai, Y., Machida, T., Shirouzu, I., et al. (2014a). Energy landscapes of resting-state brain networks. Frontiers in Neuroinformatics, 8, 12.

Watanabe, T., Masuda, N., Megumi, F., Kanai, R., Rees, G. (2014b). Energy landscape and dynamics of brain activity during human bistable perception. Nature Communications, 5, 4765.

Watanabe, T., Rees, G. (2017). Brain network dynamics in high-functioning individuals with autism. Nature Communications, 8, 16048.

Weatherby, A. J., P. H. Warren, R. Law. (1998). Coexistence and collapse: an experimental investigation of the persistent communities of a protist species pool. Journal of Animal Ecology 67, 554–566.

Weiher, E., Keddy, P. (Eds.). (2001). Ecological assembly rules: perspectives, advances, retreats. Cambridge University Press.

Widder, S., Allen, R. J., Pfeiffer, T., Curtis, T. P., Wiuf, C., Sloan, W. T., et al. (2016). Challenges in microbial ecology: building predictive understanding of community function and dynamics. The ISME journal, 10(11), 2557–2568.

Wilsey, B. J., Barber, K., Martin, L. M. (2015). Exotic grassland species have stronger priority effects than natives regardless of whether they are cultivated or wild genotypes. New Phytologist, 205, 928–37.

Wilson, W. G., Deroos, A. M., McCauley, E. (1993). Spatial instabilities within the diffusive Lotka-Volterra system: individual-based simulation results. Theoretical population biology, 43(1), 91–127.

Wootton, J. T. (2001). Prediction in complex communities: analysis of empirically derived Markov models. Ecology, 82(2), 580–598.

Young, T. P., Petersen, D. A., Clary, J. J. (2005). The ecology of restoration: historical links, emerging issues and unexplored realms. Ecology letters, 8, 662–73.

Young, T. P., Zefferman, E. P., Vaughn, K. J., Fick, S. (2015). Initial success of native grasses is contingent on multiple interactions among exotic grass competition, temporal priority, rainfall, and site effects. AoB PLANTS, 7, 081.

## References

1. Bickel, P., K. Doksum. (1977). Mathematical Statistics: Basic Ideas and Selected Topics. San Francisco: Holden-Day.

2. Azaele, S., Muneepeerakul, R., Rinaldo, A., Rodriguez-Iturbe, I. (2010). Inferring plant ecosystem organization from species occurrences. Journal of theoretical biology, 262(2), 323–329.

3. Murphy, K. P. (2012). Machine Learning: A Probabilistic Perspective. The MIT Press.

4. Harris, D. J. (2015). Multi-Process Statistical Modeling of Species’ Joint Distributions. University of California, Davis.

5. Lee, J. D., Hastie, T. J. (2015). Learning the structure of mixed graphical models. Journal of Computational and Graphical Statistics, 24(1), 230–253.

6. Watanabe, T., Hirose, S., Wada, H., Imai, Y., Machida, T., Shirouzu, I., et al. (2014a). Energy landscapes of resting-state brain networks. Frontiers in Neuroinformatics, 8, 12.

7. Watanabe, T., Masuda, N., Megumi, F., Kanai, R., Rees, G. (2014b). Energy landscape and dynamics of brain activity during human bistable perception. Nature Communications, 5, 4765.

8. Harris, D. J. (2016). Inferring species interactions from co-occurrence data with Markov networks. Ecology, 97(12), 3308–3314.

9. Robbins, H., S. Monro. 1951. A Stochastic Approximation Method. The Annals of Mathematical Statistics 22:400–407.

10. Salakhutdinov, R., and G. Hinton. (2012). An efficient learning procedure for deep Boltzmann machines. Neural Computation 24:1967–2006.

11. Younes, L. 1999. On the convergence of Markovian stochastic algorithms with rapidly de-creasing ergodicity rates. Stochastics: An International Journal of Probability and Stochastic Processes 65:177–228.

12. Hinton, G. E. (2012). A practical guide to training restricted Boltzmann machines. In Neural networks: Tricks of the trade (pp. 599–619). Springer, Berlin, Heidelberg.

